# Optimal compensation for neuron death

**DOI:** 10.1101/029512

**Authors:** David G.T. Barrett, Sophie Denève, Christian K. Machens

## Abstract

The brain has an impressive ability to withstand neural damage. Diseases that kill neurons can go unnoticed for years, and incomplete brain lesions or silencing of neurons often fail to produce any effect. How does the brain compensate for such damage, and what are the limits of this compensation? We propose that neural circuits optimally compensate for neuron death, thereby preserving their function as much as possible. We show that this compensation can explain changes in tuning curves induced by neuron silencing across a variety of systems, including the primary visual cortex. We find that optimal compensation can be implemented through the dynamics of networks with a tight balance of excitation and inhibition, without requiring synaptic plasticity. The limits of this compensatory mechanism are reached when excitation and inhibition become unbalanced, thereby demarcating a recovery boundary, where signal representation fails and where diseases may become symptomatic.

## Introduction

The impact of neuron loss on information processing is poorly understood (Montague et al., 2012; Palop et al., 2006). Every day, approximately 85,000 neurons die in the healthy adult brain (Morrison and Hof, 1997; Pakkenberg and Gundersen, 1997). Chronic diseases such as Alzheimer’s disease and brain tumor growth increase this death rate considerably, and acute events such as a stroke and traumatic brain injury can kill huge numbers of cells rapidly. Yet, this damage can often be asymptomatic and go unnoticed for years (Leary and Saver, 2003). Similarly, the incomplete lesion or silencing of a targeted brain area may ‘fail’ to produce any measurable, behavioral effect. This resilience of neural systems to damage is especially impressive when compared to man-made computer systems that typically lose all function following only minor destruction of their circuits. A thorough understanding of the interplay between neural damage and information processing is therefore crucial for our understanding of the nervous system, and may also help in the interpretation of various experimental manipulations such as pharmacological silencing (Aksay et al., 2007; Crook and Eysel, 1992), lesion studies (Keck et al., 2008), and optogenetic perturbations (Fenno et al., 2011).

In contrast to the study of information representation in damaged brains, there has been substantial progress in our understanding of information representation in healthy brains (Abbott, 2008). In particular, the theory of efficient coding, which states that neural circuits represent sensory signals optimally given various constraints (Barlow, 1961; Olshausen and Field, 1996; Olshausen and Simoncelli, 2001; Rieke et al., 1997; Salinas, 2006), has successfully accounted for a broad range of observations in a variety of sensory systems, in both vertebrates (Greene et al., 2009; Olshausen and Field, 1996; Olshausen and Simoncelli, 2001; Smith and Lewicki, 2006) and invertebrates (Fairhall et al., 2001; Machens et al., 2005; Rieke et al., 1997). However, an efficient representation of information is of little use if it cannot withstand some damage, such as normal cell death. Plausible mechanistic models of neural computation should be able to withstand the type of damage that the brain can withstand.

In this work, we propose that neural systems maintain stable signal representations by actively compensating for the destruction of neurons. We show that this compensation does not require synaptic plasticity, but can be implemented instantaneously by a balanced network whose dynamics and connectivity are tuned to implement efficient coding (Boerlin et al., 2013; Bourdoukan et al., 2012). When too many cells die, this balance is disrupted, and the signal representation is lost. We predict how much cell death can be tolerated by a neural system and how tuning curves change shape following optimal compensation. We illustrate these predictions using three specific neural systems for which experimental data before and after silencing are available - the oculomotor integrator in the hindbrain (Aksay et al., 2000, 2007), the cricket cercal system (Mizrahi and Libersat, 1997; Theunissen and Miller, 1991), and the primary visual cortex (Crook and Eysel, 1992; Crook et al., 1996, 1997, 1998; Hubel and Wiesel, 1962). In addition, we show that many input/output non-linearities in the tuning of neurons can be re-interpreted to be the result of compensation mechanisms within neural circuits. Therefore, beyond dealing with neuronal death, the proposed optimal compensation principle expands the theory of efficient coding and provides important insights and constraints for any neural code.

## Results

**The principle of optimal compensation.** We begin our illustration of optimal compensation using a simple two-neuron system that represents a signal, *x*(*t*). This signal may be a time-dependent sensory signal such as luminosity for example, or more generally, may be the result of a computation from within a neural circuit. We make two assumptions about how this signal is represented. First, we assume that an estimate of the signal can be extracted from the neurons by summing their instantaneous firing rates:

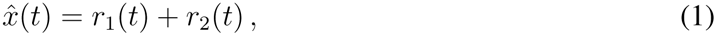

where *r*_1_(*t*) is the firing rate of neuron 1, *r*_2_(*t*) is the firing rate of neuron 2, and 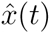 is the signal estimate. Second, we assume that the representation performance can be quantified using a simple cost-benefit trade-off. This trade-off is called our loss function and is given by:

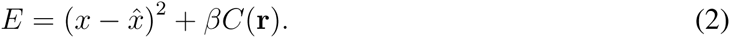

Here, the first term quantifies the signal representation error - the smaller the difference between the readout 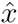 and the actual signal *x*, the smaller the representation error. The second term quantifies the cost of the representation and is given by 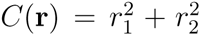. This term acts as a regularizer, ensuring that signal representations are shared amongst all neurons using low firing rates. The parameter *β* determines the trade off between this cost and the representation error. Quadratic loss functions such as Equation 2 have been a mainstay of efficient coding theories, for instance in the context of stimulus reconstruction (Rieke et al., 1997) or sparse coding (Olshausen and Field, 1996).

We can now study how firing rates must change in this system to compensate for the death of a neuron. In the healthy, undamaged state, there are many possible firing rate combinations that can represent a given signal (Figure 1A, dashed line). Among these combinations, the solution with equal firing rates is optimal, because it has the smallest cost (Figure 1A, black circle). Now, we kill neuron 1 in our simple model, and ask: how should the firing rate of neuron 2 change, in order to maintain the representation performance? We see that its firing rate will double (for very small *β*) so as to compensate for the failure of neuron 1 (Figure 1A, red circle). We obtain this result by minimizing the loss function (Equation 2) under the constraint that *r*_2_ = 0. This constraint is the mathematical equivalent of neural death. The induced change in the firing rate of neuron 2 preserves the signal representation, and thereby compensates for the loss of neuron 1.

**Figure 1.**
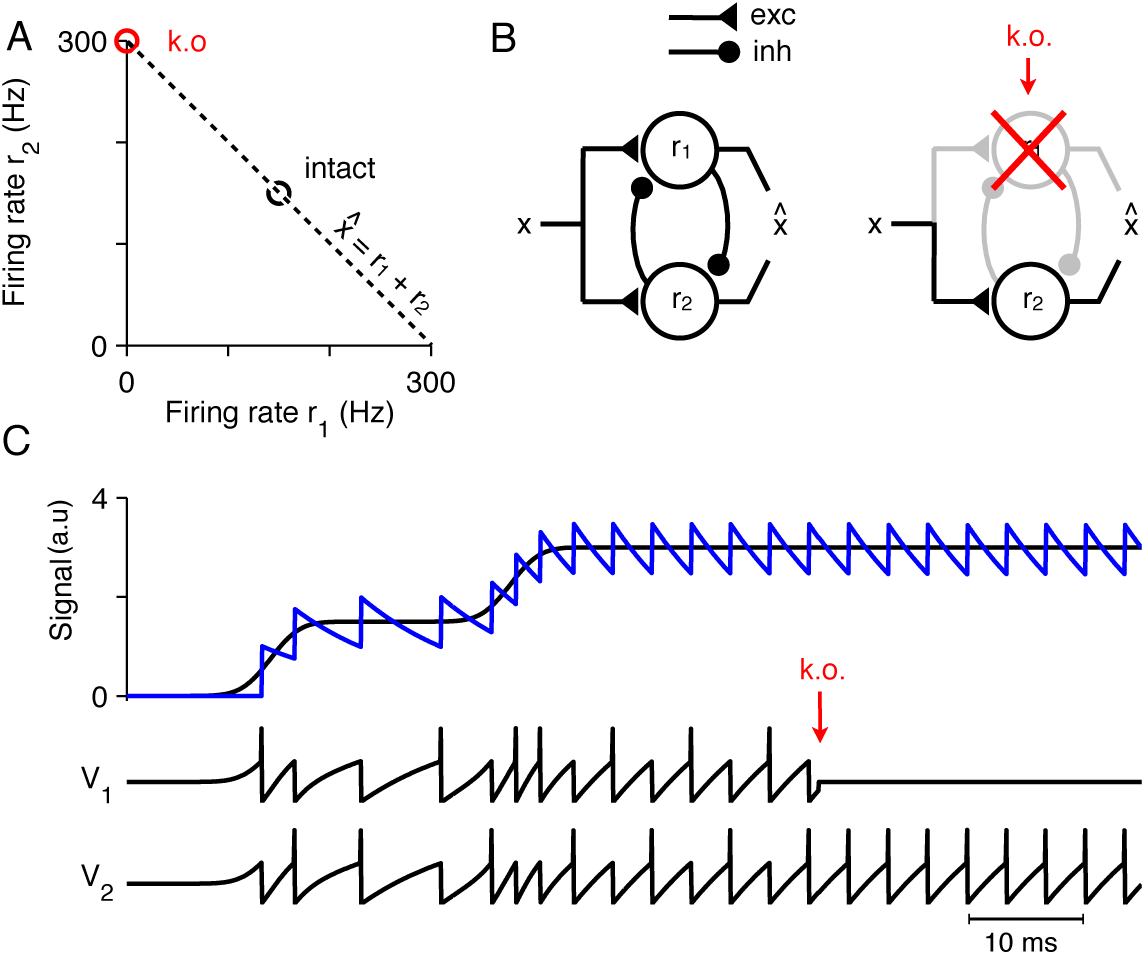
Optimal representation and optimal compensation in a two-neuron example. **(A)** A signal *x* can be repre using many different firing rate combinations (dashed line). Before cell death, the combination that best shares the load between the neurons, requires that both cells are equally active (black circle). After the death of neuron 1, the signal must be represented entirely by the remaining neuron, and so, its firing rate must double (red circle). This is the optimal compensation, for this two-neuron example. (**B**) This optimal compensation can be implemented in a network of neurons that are coupled by mutual inhibition and driven by an excitatory input signal. When neuron 1 is knocked out, the inhibition disappears, allowing the firing rate of neuron 2 to increase. (**C**) Two spiking neurons, connected together as in **B**, representing a signal *x* (black line, top panel) by producing appropriate spike trains and voltage traces, *V*_1_ and *V*_2_ (lower panels). To read out the original signal, the spikes are replaced by decaying exponential (a simple model of postsynaptic potentials), and summed to produced a readout, 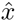 (blue, top panel). After the death of neuron 1 (red arrow), neuron 2 compensates by firing twice as many spikes.

The system can compensate the loss of a neuron because it is redundant (two neurons for one signal). While such redundancy is a necessary prerequisite for robustness, our toy example shows that it is by no means sufficient: to keep functionality, and to properly represent the signal, the firing rate of the remaining neuron needs to change.

**Optimal compensation through instantaneous restoration of balance.** Next, we investigate how this compensation can be implemented in a neural circuit. One possibility is that a circuit rewires through internal plasticity mechanisms in order to correct for the effect of the lost neurons. Surprisingly, we find that plasticity is not necessary. Rather, neural networks can be wired up such that their internal dynamics moves the instantaneous firing rates rapidly into the minimum of the loss function (Equation 2) (Boerlin et al., 2013; Charles et al., 2012; Hu et al., 2012; Rozell et al., 2008). As we will show, such dynamics are sufficient to compensate for the loss of neurons.

First, we assume that instantaneous firing rates, *r_i_*(*t*), are equivalent to filtered spike trains, similar to the postsynaptic filtering of actual spike trains (see Methods for details). Specifically, a spike fired by neuron *i* contributes a discrete unit to its instantaneous firing rate, *r_i_* → *r_i_* + 1, followed by an exponential decay. Second, we assume that each neuron will only fire a spike if the resulting change in the instantaneous firing rate reduces the loss (Equation 2). From these two assumptions, the dynamics and connectivity of the network can be derived (Boerlin et al., 2013; Bourdoukan et al., 2012). Specifically, we obtain a network containing two integrate-and-fire neurons driven by an excitatory input signal and coupled by mutual inhibition (Figure 1B). Both neurons work together, and take turns at producing spikes (Figure 1C). The neuron that reaches threshold first produces a spike, contributes to the signal readout and inhibits the other neuron. Now, when a neuron dies (Figure 1C, red arrow), the remaining neuron no longer receives inhibition from its partner neuron and it becomes much more strongly excited, spiking twice as often. In other words, the remaining neuron now implements the full optimization alone. This compensation happens because each neuron seeks to minimize the loss function, with or without the help of other neurons. When one neuron dies, the remaining neuron automatically and rapidly assumes the full burden of signal representation. In this way, the simple spiking network naturally implements optimal compensation following cell death (see Methods for additional details).

These ideas can be extended to realistic neural circuits by scaling up to larger networks. In a network of *N* neurons, the signal is represented by the weighted summation of *N* firing rates:

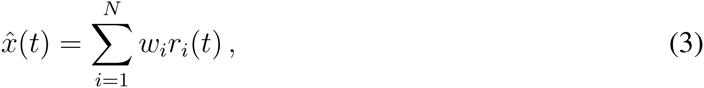

where *w_i_* is the readout weight of the *i^th^* neuron and *r_i_*(*t*) is the instantaneous firing rate of the *i^th^* neuron (Figure 2A–C). Such linear readouts are used in many contexts and broadly capture the integrative nature of dendritic summation. Just as for the two-neuron network, we can then derive both network connectivity and dynamics for this larger network, by assuming that its spiking dynamics minimize the loss function (Equation 2) (Boerlin et al., 2013; Bourdoukan et al., 2012).

**Figure 2.**
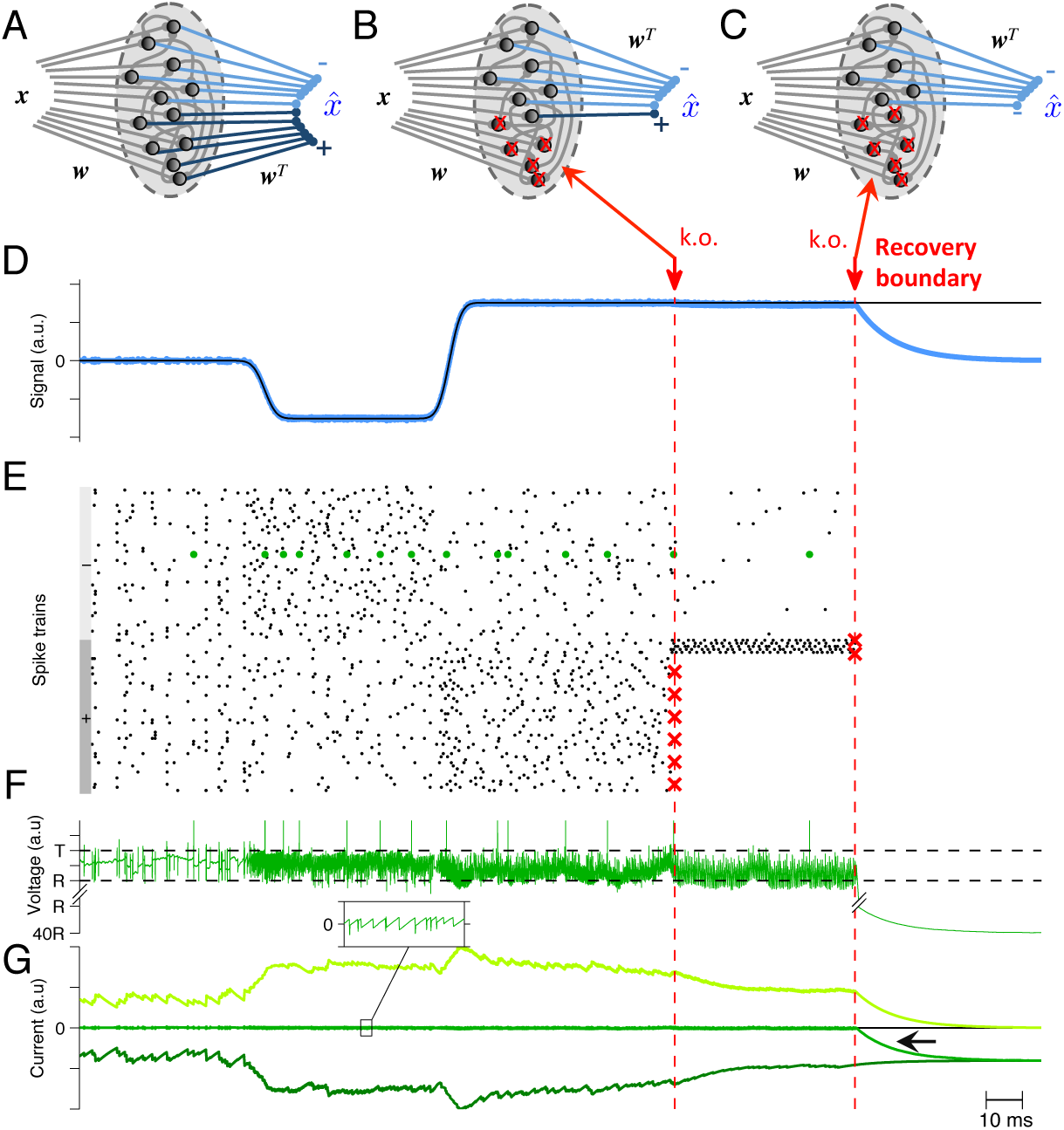
Optimal compensation and recovery boundary in a larger spiking network (*N* = 100 neurons) **(A)** Schematic of a network representing *x*(*t*). The signal *x*(*t*) can be read out by post-synaptically filtering each spike train, multi each filtered spike train with its readout weights m_;_, and then summing them all to produce 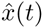. The readout weight for each neuron is either positive (dark blue) or negative (bright blue). (**B**) Schematic of a network where a large portion of neurons with positive readout weights are knocked out. (**C**) Schematic of a network where 100% of neurons with positive readout weights are knocked out. (**D**) The network provides an accurate representation 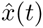 (blue line) of the time-varying signal *x* (black line). When 90% of neurons with positive readout weights are knocked out (k.o.), firing rates in the remaining neurons change to compensate the loss. However, when 100% of neurons with positive readout weights are knocked out, the representation accuracy degrades (k.o. at the recovery boundary). **(E)** A raster plot of spiking activity shows that spike trains are quite irregular. Neurons with positive and negative readout weights are marked by grey bars on the left. Red crosses mark the portion of neurons that are knocked out. Whenever neurons are knocked out, the remaining neurons change their firing rates instantaneously to compensate. **(F)** Voltage of an example neuron (marked with green dots in **E)** with threshold *T* and reset potential *R.* **(G)** The total excita (light green) and inhibitory (dark green) input currents to this example neuron are balanced, and the remaining fluctuations (inset) produce the membrane potential shown in **F.** This balance is maintained after partial knock out of neurons. However, when all the neurons with positive readout weights die, the network becomes unbalanced (black arrow), because there is not enough excitatory current (light green) to balance the inhibitory currents (dark green). This is the recovery boundary for the network (see also Figure S1).

The resulting network consists of leaky integrate-and-fire neurons receiving a tight balance of excitatory and inhibitory inputs (Figure 2D–G). In this balanced state, the signal *x*(*t*) is represented accurately (Figure 2D). Now, when some portion of the neural population dies, the remaining neurons receive less inhibition (or excitation, depending on which neurons die). In turn, their firing rates change, and the overall levels of inhibition (or excitation) readjust (Figure 2G), until balance is restored, and with it, accurate signal representation is restored. In this way, tightly balanced network dynamics compensate automatically and rapidly for the destruction of their elements (see Methods for additional details).

**The recovery boundary.** As expected, this recovery from neural death is not unlimited. When too many cells die, some portions of the signal can no longer be represented, no matter how the remaining neurons change their firing rates (Figure 2D–G). The resultant *recovery boundary* coincides with a breakdown in the balance of excitation and inhibition - where there are no longer enough inhibitory cells to balance excitation (or enough excitatory cells to balance inhibition) (Figure 2G). In this example, the recovery boundary occurs when all the neurons with positive valued readout weights have died, so that the network can no-longer represent positive valued signals or remain balanced (Figure 2G, black arrow).

When all but one neuron with positive readout weights die, the remaining neuron will shoulder the full representational load. This is an unwieldy situation because the firing rate of this neuron must become unrealistically large to compensate for all the dead neurons. In a more realistic scenario, neural activity can saturate, in which case the recovery boundary occurs much earlier (see Methods and Figures S1A, B). For instance, we find that a system with an equal number of positive and negative readout weights tolerates up to 90% neuron loss (when neurons are killed in random order) if the maximum firing rate of a neuron is 1000 Hz, whereas it will only tolerate 50% neuron loss if the maximum firing rate is 150 Hz (Figure S1B).

**Analyzing tuning curves using quadratic programming.** A crucial signature of the optimal compensation principle introduced above is that neuron firing rates must change to keep the linear readout of the signal *x* stable. In order to test how these changes compare with experimental recordings, we need to quantify them more precisely, which we will do by deriving a direct mathematical relationship between the firing rates in our spiking network and the input signal.

While the relation between the dynamics of a spiking network and the observed firing rates of individual neurons has traditionally been a hard problem, addressed for instance in the framework of mean-field theory (Renart et al., 2003), we can here circumvent many of these difficulties by noting that our spiking network minimizes the loss function (Equation 2). While this minimization has been performed for instantaneous firing rates, *r_i_*(*t*), and time-varying signals *x*(*t*), we conjecture that the *average* firing rate, 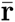, for a *constant* input signal, *x*, must be given by the minimum of the same loss function as in Equation 2 so that,

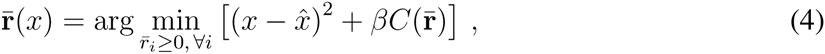

where the signal estimate, 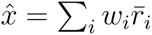, is formed using the average firing rates. This minimization is performed under the constraint that firing rates must be positive, since the firing rates of spiking neurons are positive valued quantities, by definition. Traditionally, studies of population coding or efficient coding usually assume both positive and negative firing rates (Dayan and Abbott, 2001; Olshausen and Field, 1996; Olshausen and Simoncelli, 2001), and the restriction to positive firing rates is generally considered an implementational problem (Rozell et al., 2008). Surprisingly, however, we find that the constraint changes the nature of the solutions and provides fundamental insights into the shapes of tuning curves in our networks, as clarified below.

Mathematically, the minimization of a quadratic function under linear constraints is called *quadratic programming.* In our case, the loss function is quadratic in the average firing rates, 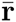, and the positivity constraint is linear in the firing rates. Using quadratic programming, we can therefore calculate the average firing rates 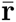 by solving Equation 4 for a range of values of *x*. This procedure produces neural tuning curves, 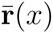 (Figure 3, second column), that closely match those measured in simulations of our tightly balanced network (Figure 3, third and fourth column), for a variety of networks of increasing complexity (Figure 3; see supplementary materials for mathematical details).

**Figure 3.**
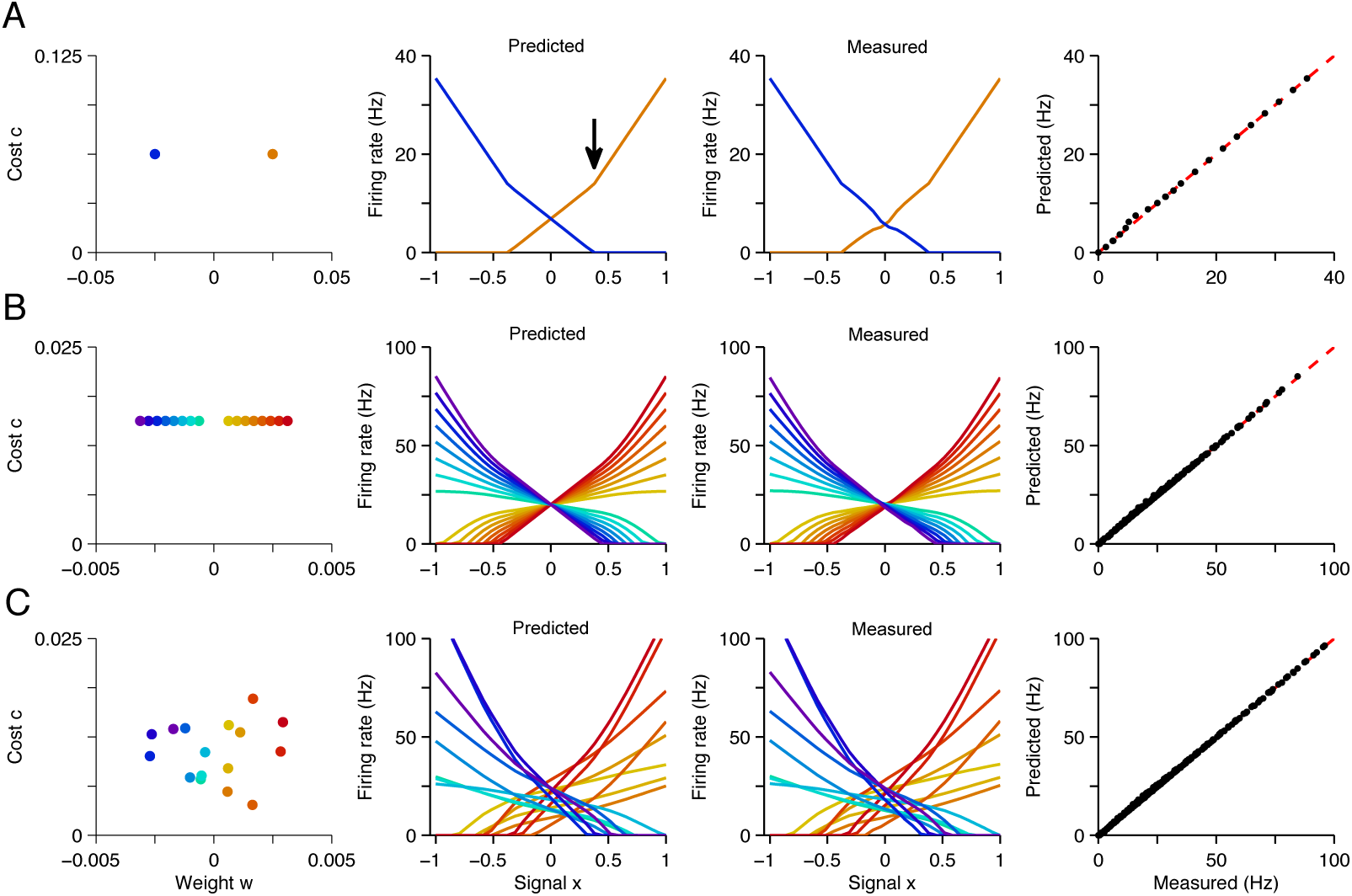
Explaining tuning curves in the spiking network with quadratic programming. **(A)** A network with two neurons. The first column shows the decoding weights {*w*_1_, *w*_2_} and the cost terms for each neuron {*c*_1_, *c*_2_}. The second column shows the tuning curves predicted using quadratic programming. The third column shows the tuning curves measured during a 1 second simulation of the respective spiking network. The fourth column shows the match between the measured and predicted tuning curves. **(B)** Similar to **A**, but using a network of 16 neurons with inhomogeneous, regularly spaced decoding weights {*w_i_*}. The resulting tuning curves are regularly spaced. Neurons with small decoding weight magnitude have smaller thresholds and shallower slopes; neurons with large decoding weight magnitudes have larger thresholds and steeper slopes. **(C)** Similar to **B** except that decoding weights and cost terms are regularly spaced with the addition of some noise. This irregularity leads to inhomogeneity in the balanced network tuning curves, and in the quadratic programming prediction (see also Figures S2-S4).

Our first observation is that the positivity constraint produces non-linearities in neural tuning curves. We illustrate this using a two-neuron system with two opposite valued readout weights, *w*_1_ = 0.025, *w*_2_ = −0.025 (Figure 3A, first column). At signal value *x* = 0, both neurons fire at equal rates so that the readout, Equation 3, correctly becomes 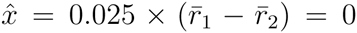. When we move to higher values of *x*, the firing rate of the first neuron, 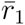, increases linearly (Figure 3A, second column, orange line), and the firing rate of the second neuron, 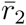, decreases linearly (Figure 3A, second column, blue line), so that in each case, 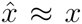. Eventually, around *x* = 0.4, the firing rate of neuron 2 hits zero (Figure 3A, black arrow). Since it cannot decrease below zero, and because the estimated value 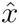 must keep growing with *x*, the firing rate of neuron 1 must grow at a faster rate. This causes a kink in its tuning curve (Figure 3A, black arrow). This kink in the tuning curve slope is an indirect form of optimal compensation, where the network is compensating for the temporary silencing of a neuron. We provide further geometric insights into the shape of the tuning curves obtained from quadratic programming, in Figure S2 and in the Methods.

Our second observation is that tuning curves depend on the decoder weight values: a neuron with a negative weight has a negative firing rate slope and a neuron with a positive weight has a positive firing rate slope (Figure 3A–C, second column). If the network consists of several neurons, then those that have small readout weight magnitude have shallow slopes and thresholds far from *x* = 0, and those with large readout weight magnitude have steep slopes and thresholds close to *x* = 0 (Figure 3B, second column). By increasing the range of readout weight values, we increase the range or tuning curve slopes (Figure S3A). Every time one of the neurons hits the zero firing rate lower bound, the tuning curve slopes of all the other neurons change.

Our third observation is that tuning curves depend on the form of the cost terms, 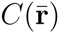. If there is no cost term, there are many equally favorable firing rate combinations that produce identical readouts (Figure 1A). The precise choice of a cost term largely determines which of these solutions is found by the network. In Figure 3, we choose a *biased* quadratic cost 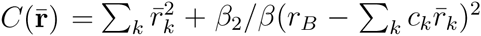. Here, the second term biases the population towards a solution where a background task variable *r_B_* is represented, and where the contribution of neuron *k* to this background task is determine by *c_k_*. This background task could be any signal-independent variable, that is not being probed or varied during an experiment. It has the effect of forcing tuning curves to intercept, thereby revealing much of the complexity of the quadratic programming solution as we have just observed. By increasing the range of neuron cost values {*c_k_*}, we increase the range or tuning curve intercepts (Figure S3B), and by decreasing the magnitude of cost values, we decrease the tuning curve intercept values (Figure S3C). As the heterogeneity of the cost parameters and readout weights increases, we see that the heterogeneity of tuning curve shapes increases (Figure 3C, second column). If we use a quadratic cost without a bias term 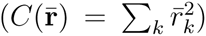, tuning curves do not intercept (Figure S4A). If we use a linear cost term 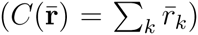, then a small number of neurons will dominate the representation (Figure S4B and Methods).

We note that although the firing rate predictions here using Equation 4 are very accurate (Figure 3, fourth column), we expect these predictions to break down in certain network regimes. For example, if membrane potential noise is large (about the same order of magnitude as the synaptic input) (Figure S5A), or if the readout weights become too large (Figure S5B and Methods). Accordingly, the dynamics of our network “implements” only an approximation to the quadratic programming algorithm, albeit a close approximation for the cases considered here. To simplify our nomenclature, we take this to be implied for our spiking network implementation, but we note that the quadratic programming solutions we obtain are the true optimal solutions.

**Tuning curves before and after neuronal silencing.** We can now calculate how tuning curves change shape in our spiking network following neuronal death. When a set of neurons are killed or silenced within a network, their firing rates are effectively set to zero. We can include this silencing of neurons in our quadratic programming algorithm by simply clamping the respective neurons’ firing rates to zero:

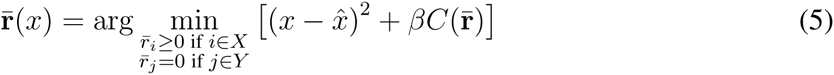

where *X* denotes the set of healthy neurons and *Y* the set of dead (or silenced) neurons. This additional clamping constraint is the mathematical equivalent of killing neurons. The difference between the solutions of Equation 4 and Equation 5 provides our definition of ‘optimal’ compensation, which is understood to be the minimization of this loss function—either numerically or through the dynamics of the spiking network. In turn, we can study the tuning curves of neurons in networks with knocked-out neurons.

As a first example, we compare the tuning curves of neurons before and after neuron death. Using a one-dimensional input signal, *x*, and similar network connectivity as before (Figure 3C), we solve Equation 4 and obtain a complex mixture of tuning curves with positive and negative slopes, and diverse threshold crossings (Figure 4A). We then calculate how these tuning curves change shape following neuron death (Figure 4B), using Equation 5. We find that neurons that have similar tuning to the knocked out neurons increase their firing rates and dissimilarly-tuned neurons decrease their firing rates. In this way, signal representation is preserved as much as possible (Figure 4B, inset). In comparison, we find that a network that does not change its tuning curves after cell death has drastically worse representation error (Figure 4B, inset). In addition, if all the neurons with positive tuning curve slopes are killed, we cross the recovery boundary, so even optimal compensation can no longer preserve signal representation (Figure 4C).

**Figure 4.**
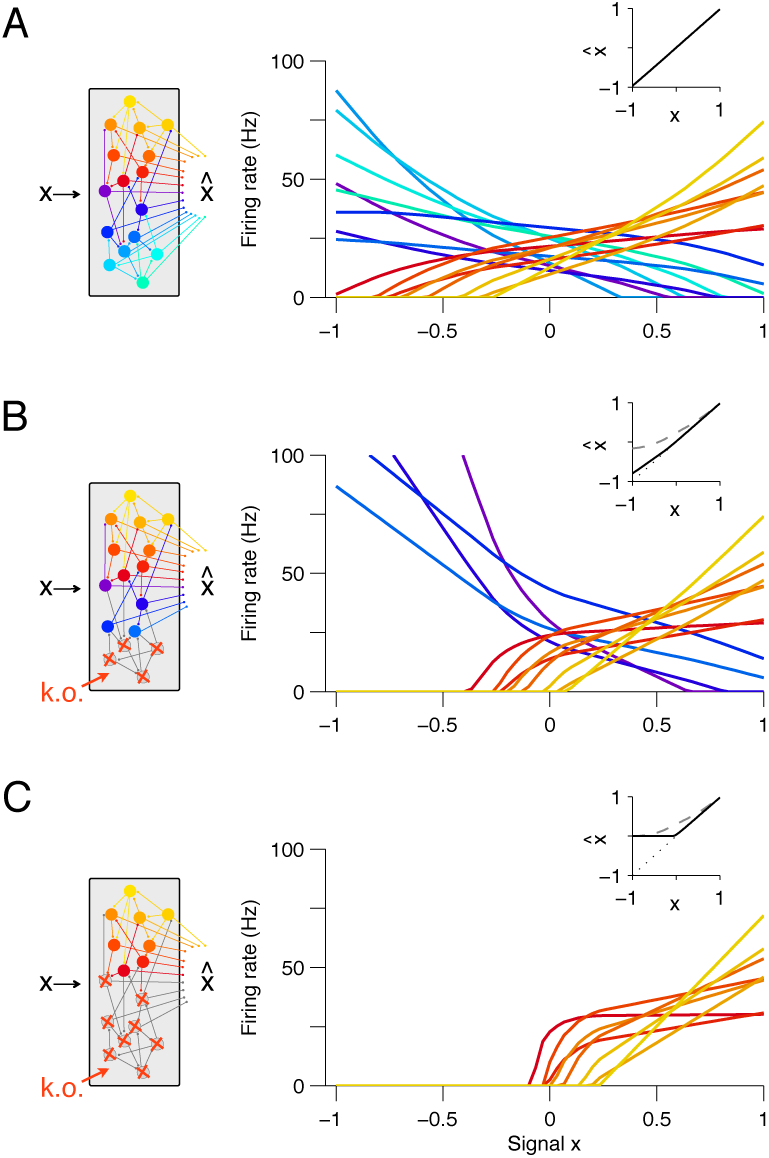
Changes in tuning curves following neuron death and optimal compensation, as calculated with quadratic programming. (**A**) The firing rates of a neural population are given as a function of input signal *x.* Here, each line is the firing rate tuning curve of a single neuron, with either positive slope (yellow–red) or negative slope (blue-green). The population contains 16 neurons (a small number, for ease of visualization). The network readout 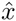 closely matches the signal *x* (inset), indicating that the network is providing a good signal representation. (**B**) Tuning curves after the death of four negatively sloped neurons and after optimal compensation from the remaining neurons. The signal representation continues to be accurate (inset, black line). If the tuning curves of the remaining neurons do not undergo optimal compensation, the signal representation fails for negative values of *x* (inset, dashed grey line). (**C**) After the death of the remaining negatively sloped neurons (bottom panel), the population is no longer capable of producing a readout 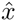 that can match negative values of the signal *x,* even with optimal compensation (inset, black line). This happens because the recovery boundary has been reached – where signal representation fails.

We note that these observations are the key signatures of optimal compensation. In particular, they are largely independent of the specific details of our network model, such as the choice of readout weights or the choice of a cost-term. For example, if instead of using a biased quadratic cost term, we use a quadratic cost term or a linear cost term, we still find that neurons with similar tuning to the knocked out neurons increase their firing rates following the knockouts (see Figure S4B, E), and we find that the recovery boundary occurs at the same place (see Figure S4C, F).

We can now make our first comparison to measurements. We consider the vertebrate oculomotor system, which is responsible for horizontal eye fixations in the hindbrain of vertebrates. This system is naturally comparable to the one-dimensional signal representation example that we have just considered: the signal *x* corresponds to the eye position, with zero representing the central eye position, positive values representing right-side eye positions and negative values representing left-side eye positions. The biased cost term could reflect the system’s desire to retain a constant muscle tone. We find that the tuning curves of neurons measured in the oculomotor system (Aksay et al., 2000) (Figure 5A) are similar to the tuning curves in our one-dimensional network calculated using Equation 4 (Figure 5B). In both cases, neurons that encode right-side eye positions have positive slopes, and these follow a recruitment order, where neurons with shallow tuning curve slopes become active before neurons with steep tuning curve slopes (Aksay et al., 2000; Fuchs et al., 1988; Pastor and Gonzalez-Forero, 2003) (Figure 5A, B inset). Now, when neurons with negatively sloped tuning curves are inactivated using lidocaine and muscimol injections (Aksay et al., 2000), the system can compensate so that it can still represent right-side eye positions (Figure 5C). This is consistent with our optimal compensation prediction (Figure 4C inset and Figure 5D). However, the system is unable to represent left-side eye positions (Figure 5C) because all the negatively sloped neurons are knocked out. This is consistent with our recovery boundary prediction (Figure 4C inset and Figure 5D).

**Figure 5.**
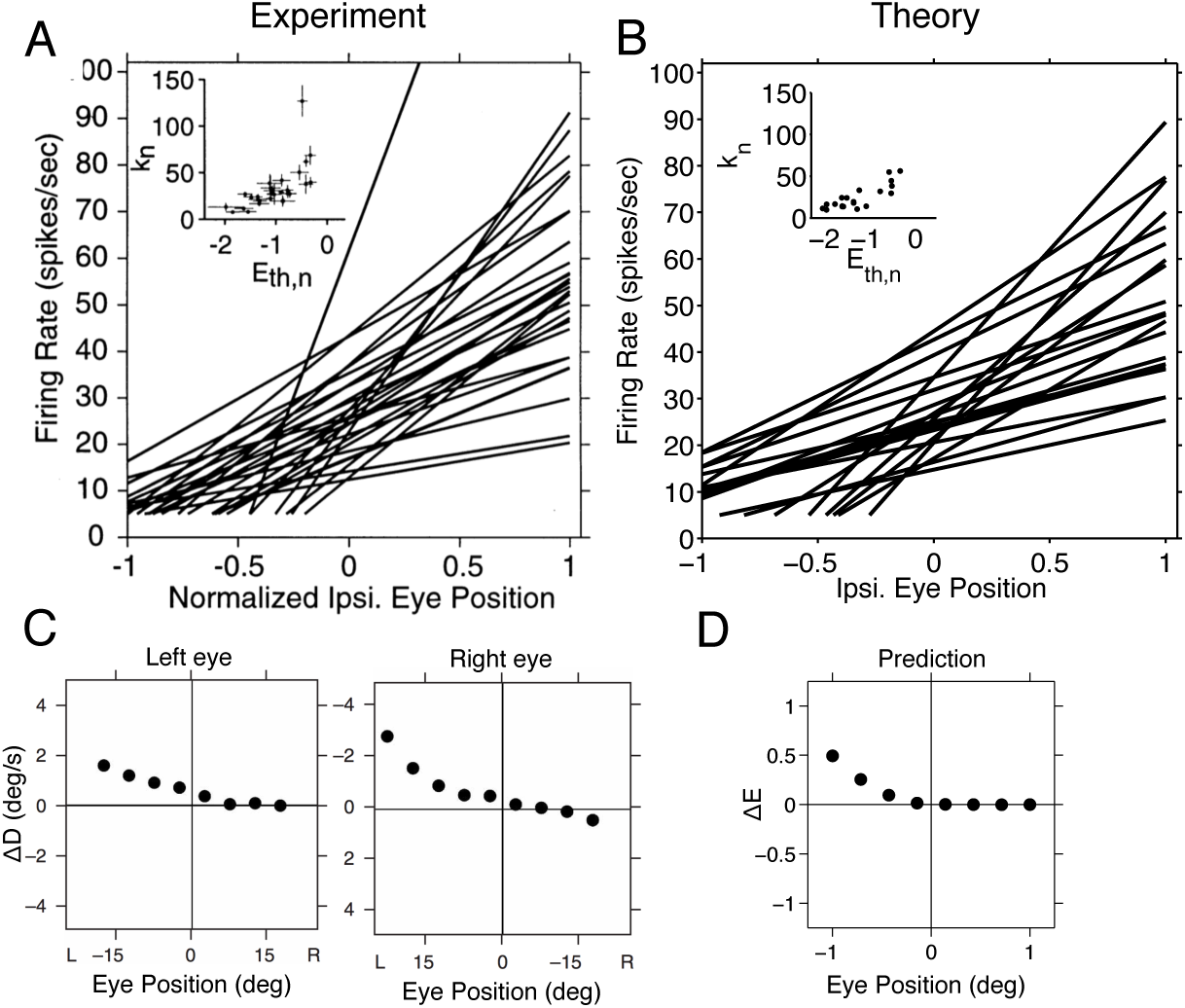
Tuning curves and inactivation experiments: Comparison between the goldfish oculomotor system and a one-dimensional (network) model. **(A)** Tuning curve measurements from *right-side* goldfish (Area 1) oculomotor neurons. Firing rate measurements above 5 Hz are fit with a straight line *f_n_ = k_n_* (*x* − *E_th_,_n_*), where *f_n_* is the firing rate of the *n^th^* neuron, *E_th_,_n_* is the firing rate threshold, *k_n_* is the firing rate slope and *x* is the eye position. As the eye position increases, from left to right, a recruitment order is observed, where neuron slopes increase as the firing rate threshold increases (inset). Reproduced from Aksay et al. (2000). **(B)** Optimal tuning curves predicted using quadratic programming (Equation 4). The predicted tuning curves are obtained using the same parameters as in previous figures (Figure 4A), and are fit using the same procedure as in the experiments from Aksay et al. (2000). As in the data, a recruitment order is observed with slopes increasing as the firing threshold increases (inset). **(C)** Eye position drift measurements after the pharmacological inactivation of left side neurons in goldfish. Inactivation was performed using lidocaine and muscimol injections. Here Δ*D = D*_after_ − *D*_before_, where *D*_before_ is the average drift in eye position before pharmacological inactivation and *D*_after_ is the average drift in eye position after pharmacological inactivation. Averages are calculated across goldfish. Adapted from Aksay et al. (2007). **(D)** Eye position representation error of a model system following optimal compensation. Here Δ*E* = *E_C_* − *E_I_*, where *E_I_* is the representation error of the intact system and *E_C_* is the representation error following optimal compensation. These representation errors are calculated using the loss function from equation 2.

**Optimal compensation in networks with bell-shaped tuning curves.** Next, we investigate these compensatory mechanisms in more complicated systems that have bell-shaped tuning curves. We consider two examples: a small sensory system called the cricket (or cockroach) cercal system, which represents wind velocity, (Theunissen and Miller, 1991) and the primary visual cortex, which represents oriented edge-like stimuli (Hubel and Wiesel, 1962). Our framework can be generalized to these cases using circular signals embedded in a two-dimensional space, x = (cos(*θ*), sin(*θ*)), where *θ* is the orientation of our signal. In the case of the cercal system, *θ* is wind-direction, and in the visual cortex, *θ* represents the orientation of an edge-like stimulus. We must also generalize our signal readout mechanism, so that our neural population can represent the two dimensions of the signal simultaneously:

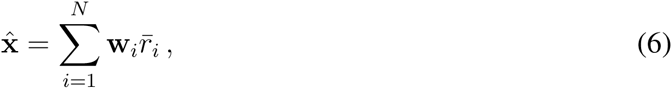

where **w***_i_* is the two-dimensional linear decoding weight of the *i^th^* neuron.

As before, we calculate tuning curves using quadratic programming (Equation 4), now for the multivariate case (Equation 28). If our system consists of four neurons with equally spaced readout weights (Figure 6A) we obtain tuning curves that are similar to the cricket (or cockroach) cercal system, with each neuron having a different, preferred wind direction (Theunissen and Miller, 1991) (Figure 6B, E). The similarity between predicted tuning curves (Figure 6B) and measured tuning curves (Figure 6E) suggests that we can interpret cercal system neurons to be optimal for representing wind direction using equally spaced readout weights.

**Figure 6.**
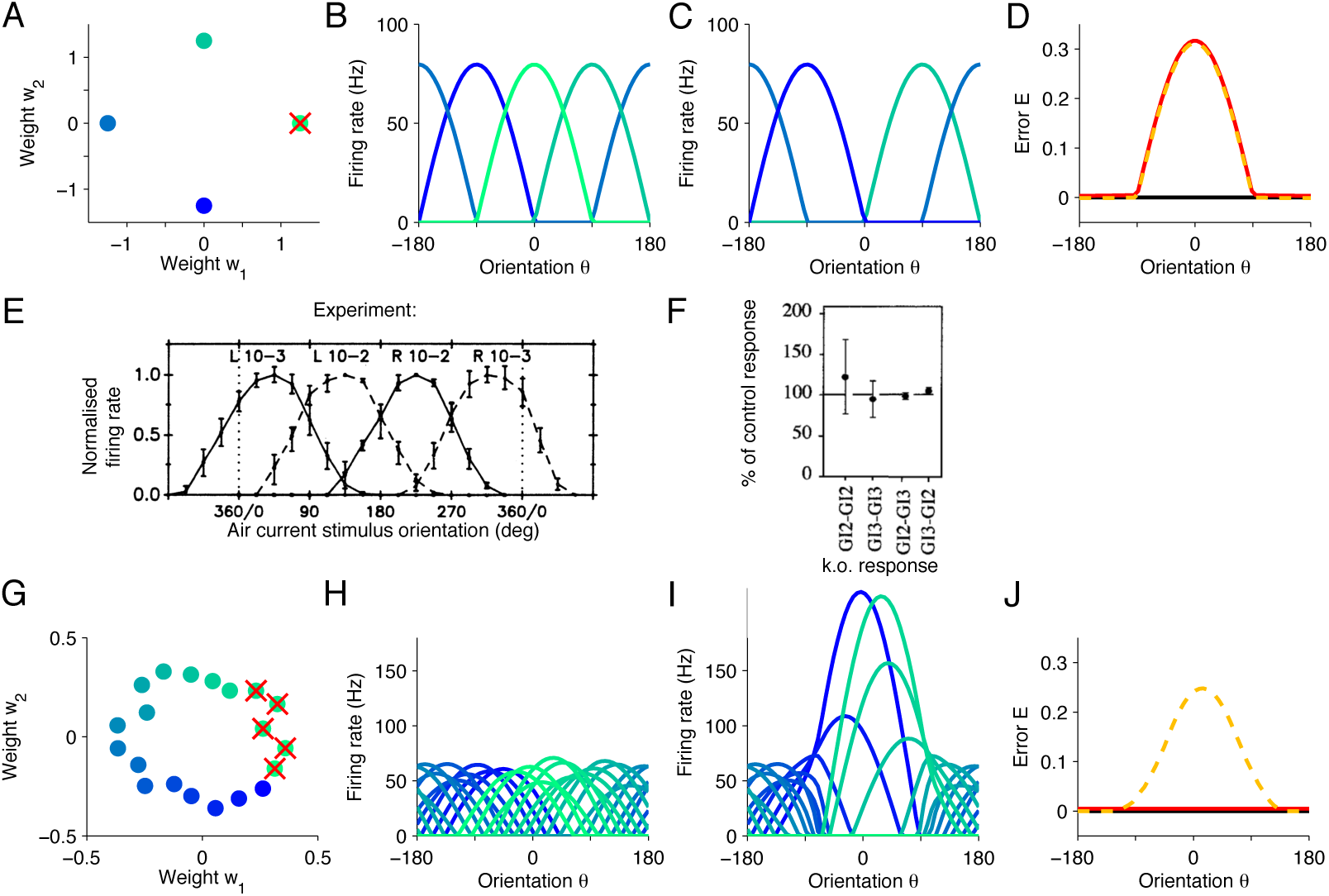
Tuning curves and inactivation experiments: Comparison between systems with angular tuning and two-dimensional (network) models. **(A)** Equally spaced readout weights for a two-dimensional network model containing four neurons. **(B)** This system has bell-shaped tuning curves that provide an optimal representation of a circular signal with orientation *θ.* **(C)** Following the death of a single neuron and optimal compensation the tuning curves of the remaining neurons do not change. **(D)** In the intact system, the representation error is negligible (black line). After cell death, the error increases strongly around the preferred orientation of the knocked out neuron (*θ* = 0*°*), with optimal compensation (red line) or without optimal compensation (dashed yellow line). **(E)** Firing rate recordings from the cricket cercal system in response to air currents at different orientations. Each neuron has a preference for a different wind direction. Compare with panel **(B).** Reproduced from Theunissen and Miller (1991). **(F)** Measured change of cockroach cercal system tuning curves following the ablation of another cercal system neuron. In all cases, there is no significant change in firing rate. This is consistent with the optimal compensation in **C.** The notation GI1-GI2 denotes that Giant Interneuron 1 is ablated and the change in Giant Interneuron 2 is measured. The firing rate after ablation is given as a percentage of the firing rate before ablation. Reproduced from Mizrahi and Libersat (1997). **(G-J)** Similar to **A-D** but for a population of 16 neurons, which creates redundancy in the representation. **(I)** When some of the neurons have been knocked out, the tuning curves of the remaining neurons change by shifting towards the preferred orientations of the knocked out neurons. **(J)** This compensation preserves high signal representation quality (red line). In comparison, if the tuning curves do not change, the readout error increases substantially (dashed yellow line).

We can now study how tuning curves in this system should change following cell death (Equation 5). This is analogous to experiments in the cockroach cercal system where a single neuron is killed (Libersat and Mizrahi, 1996; Mizrahi and Libersat, 1997). We find that our model system crosses the recovery boundary following the death of a single neuron (Figure 6D), and so, optimal compensation does not produce any changes in the remaining neurons (Figure 6C). This result is identical to measured responses (Figure 6F) (Libersat and Mizrahi, 1996; Mizrahi and Libersat, 1997). It occurs because four neurons are required, at a minimum, to represent all four quadrants of the signal space, and so, there are no changes that the remaining neurons can make to improve signal representation. Indeed, our model of the cercal system has no redundancy (see Methods for more details). As, such, the cercal system is an extreme system that exists on the edge of the recovery boundary (Libersat and Mizrahi, 1996; Mizrahi and Libersat, 1997).

The situation becomes more complicated in the large network example which has sufficient redundancy to compensate for the loss of neurons. When neurons have irregularly spaced readout weights (Figure 6G), we obtain an irregular combination of bell-shaped tuning curves (Figure 6H), similar to those found in the primary visual cortex, with each neuron having a different preferred orientation and maximum firing rate as determined by the value of the decoding weights. If we now kill neurons within a specific range of preferred orientations, then the remaining neurons increase their firing rates and shift their tuning curves towards the preferred orientations of the missing neurons (Figure 6I). In this way, the portion of space that is under-represented following cell death becomes populated by neighboring neurons, which thereby counteract the loss. The signal representation performance of the population is dramatically improved following optimal compensation compared to a system without compensatory mechanisms (Figure 6J).

**A high-dimensional example: optimal compensation in V1.** The visual cortex represents highdimensional signals, namely images, that consist of many pixel values. In the previous section, we investigated optimal compensation in a simple model of the visual cortex, in which neurons represent a one-dimensional orientation signal embedded in a two-dimensional space. Such a model is likely to be too simplistic to account for tuning curve changes in experiments. To investigate compensatory mechanisms in a more realistic model of the primary visual cortex, we make a final generalization to systems that can represent high-dimensional image signals (Figure 7A).

**Figure 7.**
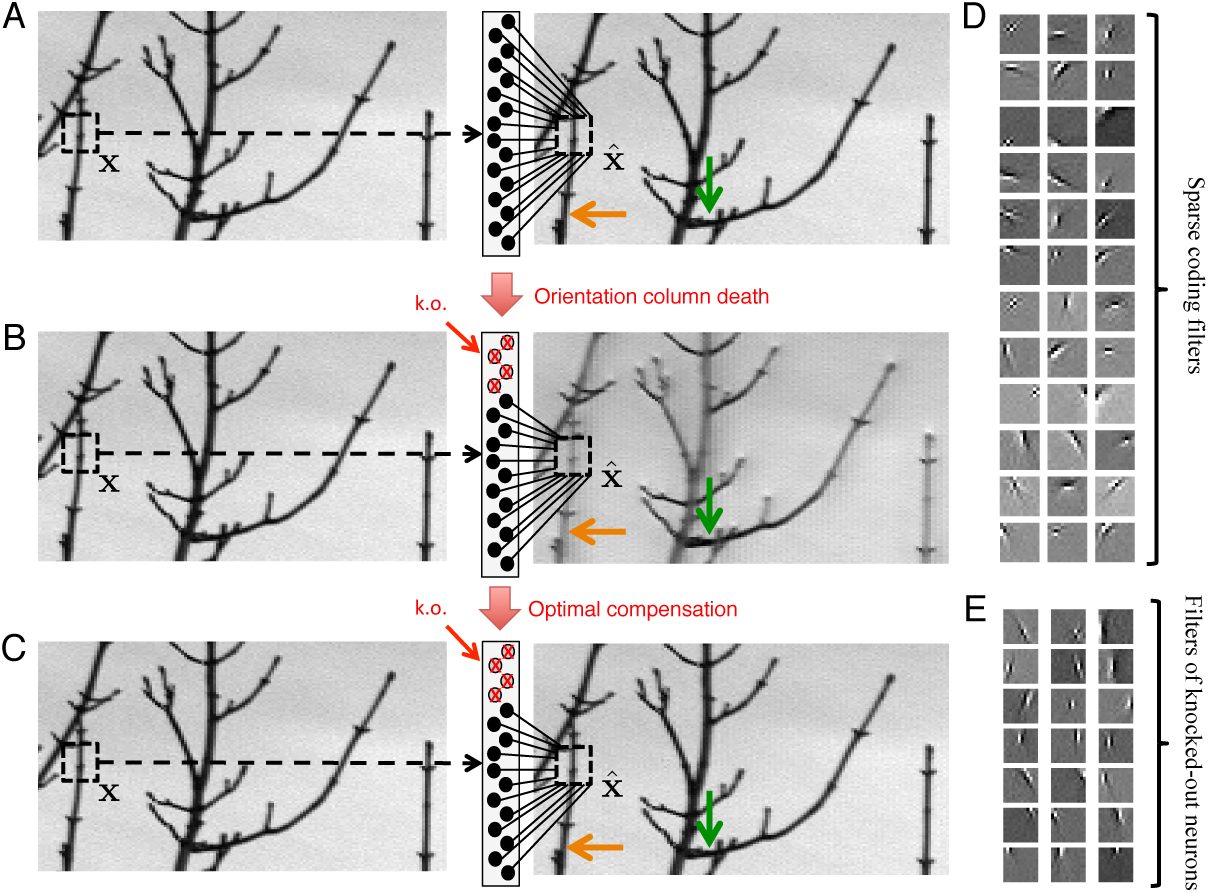
Optimal compensation for orientation column cell death in a positive sparse coding model of the visual cortex. **(A)** Schematic of a neural population (middle) providing a representation (right) of a natural image (left). This image representation is formed when neurons respond to an image patch x with a sparse representation and an output 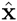. Here, patches are overlapped to remove aliasing artifacts. **(B)** Following the death of neurons that represent vertical orientations, the image representation degrades substantially without optimal compensation, especially for image segments that contain vertical lines (orange arrow), and less so for image segments that contain horizontal lines (green arrow). **(C)** Following optimal compensation, the image representation is recovered. **(D)** A selection of efficient sparse decoding weights, illustrated as image patches. Each image patch represents the decoding weights for a single neuron. In total, there are 288 neurons in the population. **(E)** A selection of vertically orientated decoding weights whose neurons are selected for simulated death. All neurons with preferred vertical orientations at 67.5*°*, 90*°* and 112.5*°* and at the opposite polarity, −67.5*°*, −90*°* and −112.5*°* are silenced.

The tuning of V1 simple cells can largely be accounted for by assuming that neural firing rates provide a sparse code of natural images (Olshausen and Field, 1996; Olshausen and Simoncelli, 2001). In accordance with this theory, we use decoding weights that are optimized for natural images, and we use a sparsity cost instead of a quadratic cost (see Methods). As before, we obtain the firing rates of our neural population by minimizing our loss function (Equation 4), under the constraint that firing rates must be positive. We find that neurons are tuned to both the orientation and polarity of edge-like images (Figure 7D), where the polarity is either a bright edge with dark flanks, or the opposite polarity - a dark edge with bright flanks. Orientation tuning emerges because natural images typically contain edges at many different orientations, and a sparse code captures these natural statistics (Olshausen and Field, 1996; Olshausen and Simoncelli, 2001). Polarity tuning emerges as a natural consequence of the positivity constraint, because a neuron with a positive firing rate cannot represent edges at two opposing polarities. Similar polarity tuning has been obtained before, but with an additional constraint that decoding weights be strictly positive (Hoyer, 2004, 2003).

Finally, we calculate optimal compensation of tuning curves following the silencing of an orientation column (Figure 7E), again using Equation 5. Without optimal compensation, (i.e., without any changes in firing rates), we find that the silencing of an orientation column damages image representation, especially for image components that contain edges parallel to the preferred orientations of the dead neurons (Figure 7B, orange arrow). When the population implements optimal compensation, the firing rates of many neurons change, and the image representation is recovered (Figure 7C). The firing rates of the remaining neurons increase substantially at the preferred orientation of the silenced cells (Figure 8A, C) and the preferred orientations of many cells shift toward the preferred orientation of the silenced cells (Figure 8A, D), similar to the simplified two-dimensional example. Note that firing rates at the anti-preferred orientation also increase (Figure 8A, C). This occurs because neurons represent more than orientation (they represent a full image), and so, neurons that have anti-preferred orientations relative to each other may have similar tuning along other signal dimensions. These compensatory firing rate changes are consistent with experiments in the cat visual cortex, in which orientation columns are silenced using GABA (Crook et al., 1996, 1997, 1998) (Figure 8B and Figure S6). Neurons that escape silencing in the cat visual cortex also shift their tuning curves towards the preferred orientations of the silenced neurons (Figure 8B). In addition, the changes observed in the cat visual cortex are rapid, occurring at the same speed as the GABA-ergic silencing, which is consistent with the rapid compensation that we predict.

**Figure 8.**
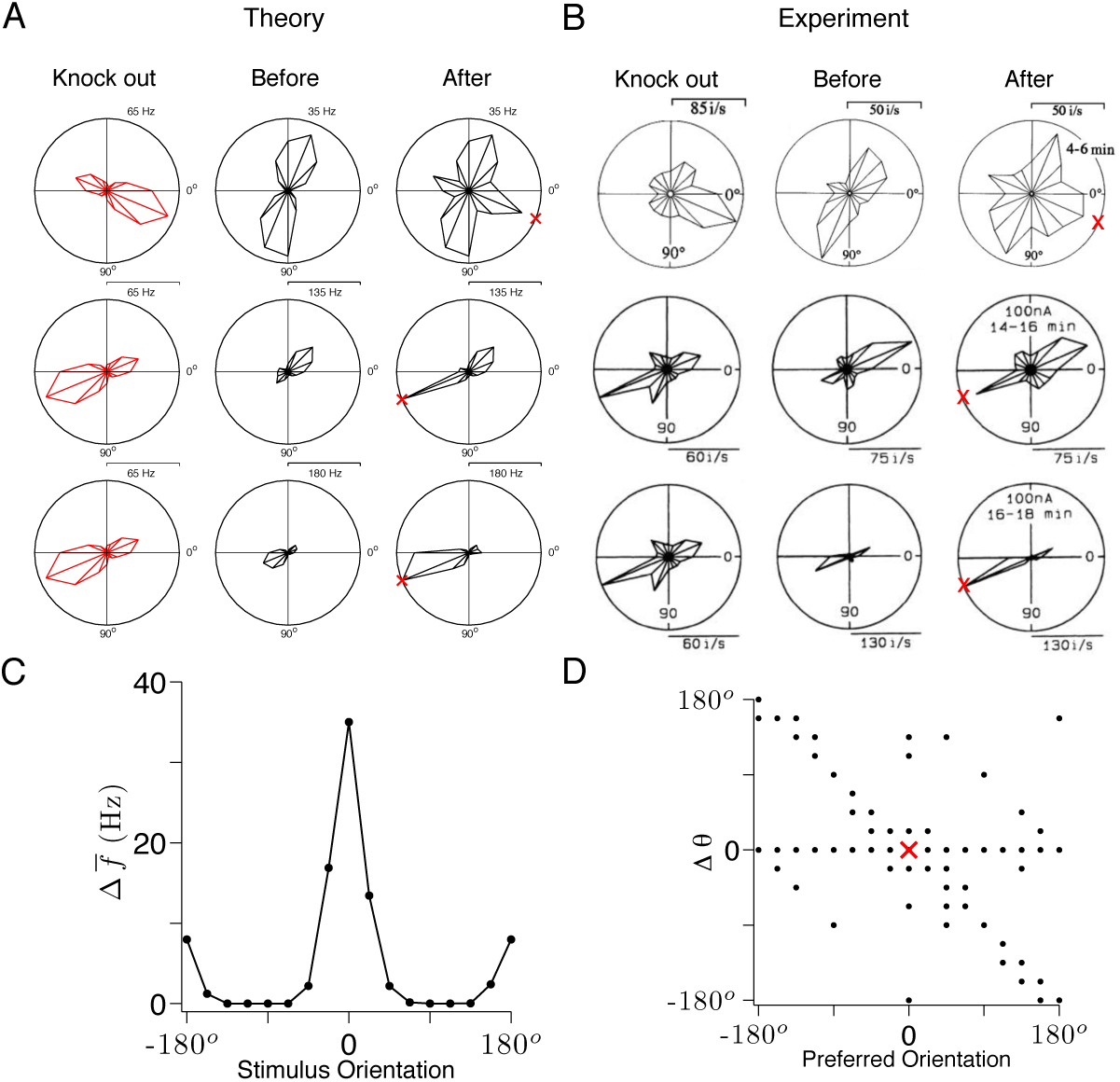
Tuning curves and inactivation experiments: Comparison between V1 recordings and the positive sparse coding model. **(A)** A set of similarly tuned neurons are selected for artificial knock out in our V1 model. The average tuning curve of these neurons is plotted in polar co-ordinates (red). The tuning curve of a test neuron is shown before the selected neurons are knocked out (middle column) and after optimal compensation (right column). Each row illustrates the response of a different test neuron to neuron death. Following optimal compensation, the neurons remaining after cell death shift their tuning curves towards the preferred orientation of the knocked out neurons (indicated by a red cross). **(B)** Recordings of cat visual cortex neurons (Area 18) both before (middle column) and after (right column) the GABA-ergic knock out of neighboring neurons (left column). The firing rates of all neurons increase in the direction of the preferred orientation of the silenced neurons (red cross). Each row illustrates the response of a different test neuron to silencing. Examples are selected for ease of comparison with **A.** Knock out measurements were obtained using multi-unit recording and single-unit recordings were obtained for neurons that were not subject to silencing. These results are adapted from Crook et al. (1996) and Crook et al. (1998). **(C)** The average change in firing rate due to optimal compensation 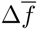 is calculated at each stimulus orientation, where the average is taken across the neural population. There is a substantial increase in firing rates close to the preferred orientations of the knocked out cells. **(D)** The change in preferred orientation Δ*θ* following optimal compensation is given for each neuron as a function of its preferred orientation before cell death. Many cells shift their preferred orientation towards the preferred orientation of the knocked out cells. Cells that do not change may still increase their firing around the preferred orientation of the knocked out neurons **(A,** top row). In **A–D** we knock out 50% of neurons with preferred orientations across a range of 50*°* (see also Figure S8).

These results are largely independent of the precise parameterization of the system, such as the number of neurons chosen and the number of neurons knocked out. A high degree of over-completeness, for instance, will shift the recovery boundary to a larger fraction of knocked-out neurons, but will not change the nature of the compensatory response (Figure S7A, B). At high degrees of over-completeness, quantitive fluctuations in tuning curve responses are averaged out, indicating that optimal compensation becomes invariant to over-completeness (Figure S7C, D). The precise proportion of neurons knocked out in the experimental studies (Crook and Eysel, 1992) is not known. However, for a range of reasonable parameter choices (50%-75% knock out within a single orientation column), our predicted tuning curve changes are consistent with the recorded population response to neuron silencing (Crook and Eysel, 1992) (Figure S6B). Although the existence of a quantitive match is interesting, it is the broad qualitative agreement between our theory and the data from a variety of systems that is most compelling.

## Discussion

To our knowledge, optimal compensation for neuron death has not been proposed before. Usually, cell death or neuron silencing is assumed to be a wholly destructive action, and the immediate neural response is assumed to be pathological, rather than corrective. Synaptic plasticity is typically given credit for recovery of neural function. For example, synaptic compensation has been proposed as a mechanism for memory recovery following synaptic deletion (Horn et al., 1996), and optimal adaptation following perturbations of sensory stimuli (Wainwright, 1999) and motor targets (Braun et al., 2009; Kording et al., 2007; Shadmehr et al., 2010) has also been observed, but on a slow timescale consistent with synaptic plasticity. In this work, we have explored the properties of optimal compensation, and have shown that it may be implemented without synaptic plasticity, on a much faster timescale, through balanced network dynamics.

The principle of optimal compensation that we have proposed here is obviously an idealization, and any putative compensatory mechanism of an actual neural system may be more limited. That said, we have catalogued a series of experiments in which pharmacologically induced tuning curve changes can be explained in terms of optimal compensation. These experiments were not originally conducted to test any specific compensatory mechansisms, and so, the results of each individual experiment were explained by separate, alternative mechanisms (Aksay et al., 2007; Crook et al., 1996, 1997, 1998; Libersat and Mizrahi, 1996; Mizrahi and Libersat, 1997). The advantage of our optimal compensation theory is that it provides a simple, unifying explanation for all these experiments.

Our work is built upon a connection between two separate theories: the theory of balanced networks, which is widely regarded to be the standard model of cortical dynamics (Renart et al., 2010; van Vreeswijk and Sompolinsky, 1996, 1998) and the theory of efficient coding, which is arguably our most influential theory of neural computation (Barlow, 1961; Bell and Sejnowski, 1997; Greene et al., 2009; Olshausen and Field, 1996; Salinas, 2006). We were able to connect these two theories for two reasons: (1) We show how to derive a tightly balanced spiking network from a quadratic loss function, simplifying the recent work of (Boerlin and Denève, 2011; Boerlin et al., 2013) by focusing on the part of their networks that generate a spike-based representation of the information. (2) We show that the firing rates in these spiking networks also obey a quadratic loss function, albeit with a positivity constraint on the firing rate. While the importance of positivity constraints has been noted in other context (Hoyer, 2004, 2003; Lee and Seung, 1999; Salinas, 2006), we here show its dramatic consequences in shaping tuning curves, which had not been appreciated previously. Now, using this connection we can easily compute the tuning curves in our network by directly minimizing the loss function under a positivity constraint. This constraint minimization problem, called quadratic programming, provides a novel link between balanced networks and traditional firing rate calculations. It allows us to think of spiking dynamics as a quadratic programming algorithm, and tuning curves as the solution. In turn, we obtain a single normative explanation for the polarity tuning of simple cells (Figure 7D), tuning curves in the oculomotor system (Figure 5A, B), and tuning curves in the cricket cercal system (Figure 6A–F), as well as the mechanisms underlying the generation of these tuning curves, and the response of these systems to neuronal loss.

Several alternative network models have been proposed that minimize similar loss functions as in our work (Charles et al., 2012; Dayan and Abbott, 2001; Hu et al., 2012; Rozell et al., 2008). However, in all these models, neurons produce positive and negative firing rates (Charles et al., 2012; Dayan and Abbott, 2001; Rozell et al., 2008), or positive and negative valued spikes (Hu et al., 2012). The compensatory response of these systems will be radically different, because oppositely tuned neurons can compensate for each other by increasing their firing rates and changing sign, whereas we predict that only similarly tuned neurons will compensate for each with increased firing rates. Similar reasoning holds for any efficient coding theories that assumes positive and negative firing rates. In more recent work, it has been found that a spiking network, that produces positive spikes only, can learn to efficiently represent natural images (King et al., 2013; Zylberberg et al., 2011). Whether these models will support optimal compensation will depend on the specifics of the learnt connectivity (see Methods). Since optimal compensation is directly related to balance, we expect any balanced network models that track signals (Renart et al., 2010; van Vreeswijk and Sompolinsky, 1996, 1998) to also implement optimal compensation following neuron death. These observations hold for generalizations of our own model in which the signal input x is replaced by slow recurrent synaptic inputs that perform more complex computations, such as arbitrary linear dynamical systems (Boerlin et al., 2013). Indeed, recent work on balanced networks that act as integrators has shown robustness to various perturbations, including neuron death (Boerlin et al., 2013; Lim and Goldman, 2013), and can therefore be considered a special case of the optimal compensation principle that we propose here.

Aside from the specific examples studied here, we can use our theory to make a number of important, testable predictions about the impact of neural damage on neural circuits in general. First, we can predict how tuning curves change shape to compensate for neuron death. Specifically, neurons with similar tuning to the dead neurons increase their firing rates and neurons with dissimilar tuning decrease their firing rates (unless they are already silent). This happens because the remaining neurons automatically seek to carry the informational load of the knocked out neurons (which is equivalent to maintaing a balance of excitation and inhibition). This is a strong prediction of our theory, and as such, an observation inconsistent with this prediction would invalidate our theory. There have been very few experiments that measure neural tuning before and after neuron silencing, but in the visual cortex, where this has been done, our predictions are consistent with experimental observations (Crook et al., 1996, 1997, 1998) (Figure 8A, B).

Our second prediction is that optimal compensation is extremely fast—faster than the timescale of neural spiking. This speed is possible because optimal compensation is supported by the balance of excitation and inhibition, which responds rapidly to neuron death—just as balanced networks can respond rapidly to changes in input (Renart et al., 2010; van Vreeswijk and Sompolinsky, 1996, 1998). We are not aware of any experiments that have explicitly tested the speed of compensation following neuron silencing. In the visual cortex, the pharmacological silencing of neurons is too slow to out-rule the possibility that there is some synaptic plasticity (Crook et al., 1996, 1997, 1998). Nonetheless, these experiments are consistent with our prediction, because the changes observed in tuning curve shape are at least as fast, if not faster than the speed of pharmacological silencing. Ideally, these predictions could be tested using neuronal ablations or optogenetic silencing.

Finally, we predict that all neural systems have a cell death recovery boundary, beyond which neural function disintegrates. Existing measurements from the oculomotor system (Aksay et al., 2007) seem to be consistent with this prediction (Figure 5 and Figure 6A–F). We predict that this recovery boundary coincides with a disruption in the balance of excitation and inhibition. This has not been explored experimentally, although the disruption of balance has recently been implicated in a range of neural disorders such as epilepsy (Bromfield, 2006) and schizophrenia (Yizhar et al., 2011). Anecdotally, there have been many unreported experiments where neural ablation has failed to cause a measurable behavioral effect. Our theory suggests that such “failed” lesion experiments may be far more interesting than previously thought, and that the boundary between a measurable and unnoticeable behavioral effect deserves specific attention. Indeed, the properties of a recovery boundary may shed light on the progression of neurodegenerative diseases—especially those that are characterized by a period of asymptomatic cell death, followed by a dramatic transition to a disabled symptomatic state, as in Alzheimer’s disease and stroke (Leary and Saver, 2003). We predict that these transitions occur at the recovery boundary of the diseased system. We also predict that an accumulation of asymptomatic damage, through aging for example, or through acute conditions such as silent stroke, will increase the susceptibility of the brain to symptomatic damage, by moving it closer to the recovery boundary. If a system was to cross our predicted recovery boundary, without there being any functional deficit, our theory would be invalidated.

These predictions, and more broadly, the principle of optimal compensation that we have developed here, promise to be useful across a number of areas. First, as a neuroscience tool, our work provides a framework for the interpretation of experimental manipulations such as pharmacological silencing (Aksay et al., 2007), lesion studies (Keck et al., 2008) and optogenetic perturbations (Fenno et al., 2011). Second, in the study of neural computation, optimal compensation may be a useful guiding principle, because plausible models of neural computation should be designed specifically to withstand the type of damage that the brain can withstand. Finally, our work provides a new framework for describing how neurodegenerative disease impacts behavior through neural networks, by generalizing the theory of efficient coding from the intact brain state to the damaged brain state (Bredesen et al., 2006; Morrison and Hof, 1997).

## Methods

We have described the properties of optimal compensation, and given a variety of examples of optimal compensation across a range of systems. Here, we present further technical details. First, we describe how we tune a spiking network to represent signals optimally and we specify our choice of parameters for each figure. For the high-dimensional sparse coding example, we describe how we calculate sparse coding receptive fields and orientation tuning curves, both before and after neuron death. Next, we explain quadratic programming with an analytically tractable example, and we provide further details on our knock-out calculations. Finally, we prove that our spiking model performs optimal compensation. The Matlab code that we used to generate all the figures in this paper, along with all the parameters used for these figures will be published online.

**Derivation of network model.** In this section, we derive the connectivity and dynamics of a network that can optimally compensate for neuron death. We consider a network of *N* leaky integrate-and-fire neurons receiving input x = (*x*_1_,&, *x_j_*,&, *x_M_*), where *M* is the dimension of the input and *x_j_* is the *j^th^* input. In response to this input, the network produces spike trains **s** = (*s*_1_,&, *s_j_*,&, *s_N_*), where 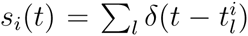 is the spike train of the *i^th^* neuron and 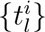 are the spike times of that neuron. Here, we describe the general formulation of our framework for arbitrary networks with *N* ≥ *M*.

A neuron fires a spike whenever its membrane potential exceeds a spiking threshold. We can write this as *V_i_* > *T_i_*, where *V_i_* is the membrane potential of neuron *i* and *T_i_* is the spiking threshold. The dynamics of the membrane potentials are given by:

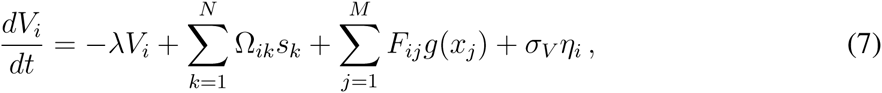

where Ω*_ik_* is the connection strength from neuron *k* to neuron *i*, *F_ij_* is the connection strength from input *j* to neuron *i*, *g*(*x_j_*) is an operator applied to the input *x_j_*, *λ* is the neuron leak and *σ_V_* is the standard deviation of intrinsic neural noise, represented by a Wiener process *η_i_* (Dayan and Abbott, 2001; Knight, 1972). The input “signal” *g*(*x_j_*) is a placeholder for any feedforward input into the networks, but also for other recurrent synaptic inputs that are not explicitly modeled here. When a neuron spikes, its membrane potential is reset to *R_i_* ≡ *T_i_* + Ω*_ii_*. Note that for ease of presentation, this reset is included in the recurrent connectivity summation of Equation 7. Also, we have written many variables without indicating their time-dependency. For example, the input signals, *x_j_*(*t*), the voltages, *V_i_*(*t*) and the spike trains, *s_k_*(*t*), are all time-dependent quantities, whereas the thresholds, *T_i_*, the leak, *λ* and the connection strengths, Ω*_ik_* and *F_ij_*, are all constants.

We assume that this network provides a representation of the input signal **x** using a simple linear decoder:

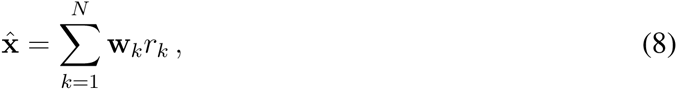

where **w***_k_* is the fixed contribution of neuron *k* to the signal. We call this a vector of readout weights or an “output kernel”. This equation is the same as Equation 6 from the main text, and is a more general version of Equation 1 and Equation 3. The instantaneous firing rate of the *k*-th neuron, *r_k_*, is a time-dependent quantity that we obtain by filtering the spike train with an exponential filter:

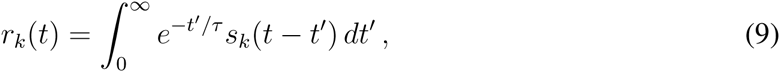

where *τ* is the time-scale of the filtering. This firing rate definition is particularly informative because it has the form of a simple model of a postsynaptic potential, which is a biologically important quantity. Note that the units of this firing rate are given by *τ*, so we must multiply by *τ*^−1^ to obtain units of Hz. For example, in our two-neuron example (Figure 1A), we want to plot *r*_1_ and *r*_2_ in units of Hz, and we have used *τ* =10 ms, so we must multiply by *τ*^−1^ = 100 Hz to obtain values in standard units.

Our goal is to tune all the parameters of this network so that it produces appropriate spike trains at appropriate times to provide an accurate representation of the input **x**, both before and after cell death. We can do this by requiring our network to obey the following rule: at a given time point *t*, each neuron only fires a spike whenever a spike reduces the loss function (Equation 2),

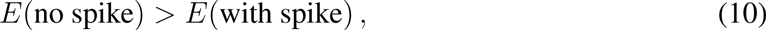

where the loss function is now assumed to be generalized to multivariate input signals **x** (Boerlin et al., 2013; Bourdoukan et al., 2012),

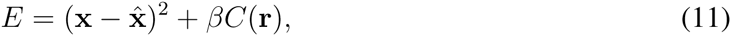

with (·)^2^ denoting an inner product, and 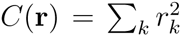. Then, since a spike changes the signal estimate by 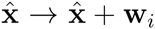 and the rate by *r_i_* → *r_i_* + 1, we can restate this spiking rule for the *i*-th neuron as:

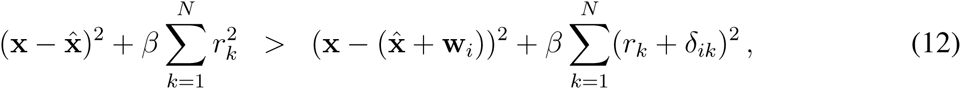

where *δ_ik_* is the kronecker delta. By expanding the right-hand-side, and canceling equal terms, this can be rewritten as

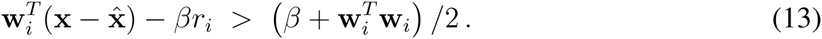

This equation describes a rule under which neurons fire to produce spikes that reduce the loss function. Since a neuron *i* spikes whenever its voltage *V_i_* exceeds its threshold *T_i_*, we can interpret the left-hand-side of this spiking condition (Equation 13) as the membrane potential of the *i^th^* neuron:

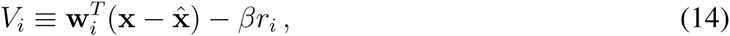

and the right-hand-side as the spiking threshold for that neuron:

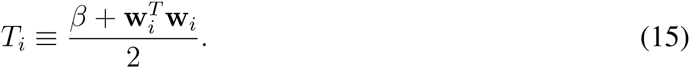

We can identify the connectivity and the parameters that produce optimal coding spike trains by calculating the derivative of the membrane potential (as interpreted in Equation 14) and matching the result to the dynamical equations of our integrate-and-fire network (Equation 7):

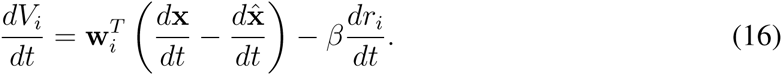

From Equation 8, we obtain 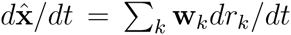 and from Equation 9 we obtain *dr_k_*/*dt* = −*r_k_*/*τ* + *s_k_*, which yields a simple differential equation for the readout:

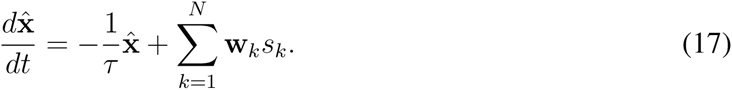

By inserting these expressions into Equation 16 we obtain:

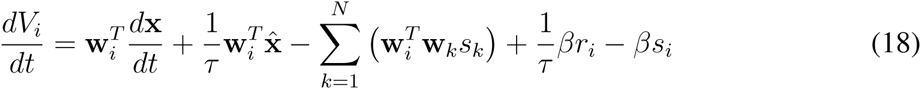

Finally, using the voltage definition from Equation 14 to write 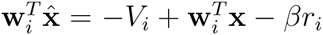 we can replace the second term on the right hand side and obtain:

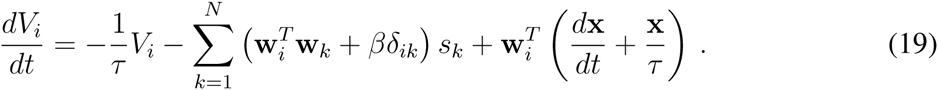

This equation describes the voltage dynamics of a neuron that produces spikes to represent signal **x**. If we now compare this equation with our original integrate-and-fire network, Equation 7, we see that these are equivalent if

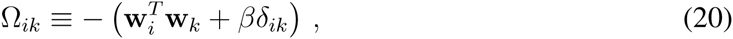

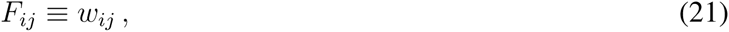

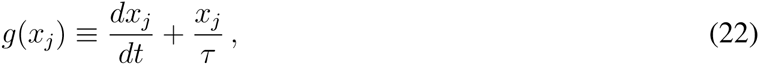

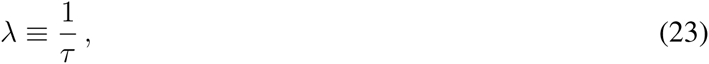

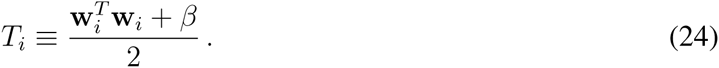

A network of integrate-and-fire neurons with these parameters and connection strengths can produce spike trains that represent the signal **x** to a high degree of accuracy. Elements of this calculation have been produced before (Boerlin et al., 2013), but are reproduced here for the sake of completeness. Also, it has been shown that this connectivity can be learned using a simple spike timing-dependent plasticity rule (Bourdoukan et al., 2012), so extensive fine-tuning is not required to obtain these spiking networks. We note that the input into the network consists of a combination of the original signal, *x_j_*, and its derivative, *dx_j_*/*dt*. In the simulations, we feed in the exact derivative. To be biologically more plausible, however, the derivative could be computed through a simple circuit that combines direct excitatory signal inputs with delayed inhibitory signal inputs (e.g. through feedforward inhibition).

Our derivation of network dynamics directly from our loss function allows us to interpret the properties of this network in terms of neural function. Each spike can be interpreted as a greedy error reduction mechanism – it moves 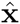 closer to the signal **x**. This error reduction is communicated back to the network through recurrent connectivity, thereby reducing the membrane potential of the other neurons. The membrane potential, in turn, can be interpreted as a representation error – a linear projection of 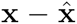 onto the neuron’s output kernel **w***_i_* (Equation 14). Following this interpretation, whenever the error becomes too big, the voltage becomes too big and it reaches threshold. This produces a spike, which reduces the error, and so on.

We can also understand these network dynamics in terms of attractor dynamics. This network implements a point attractor - firing rates evolve towards a stable fixed point in N-dimensional firing rate space. The location of this point attractor depends on neural input and network connectivity. When a neuron dies, the point attractor is projected into a subspace given by *r_k_* = 0, where neuron *k* is the neuron that has died.

Note that in this derivation, we used a quadratic cost. This cost increases the value of the spiking threshold (Equation 24) and the spiking reset (Equation 20). We can also derive network parameters for alternative cost term choices. For example, if we we use a linear cost, we simply need to drop the second term (*βδ_ik_*) in Equation 20, while keeping all other parameters the same. In other words, we can implement a quadratic cost by increasing the spiking threshold and the spiking reset, and we can implement a linear cost by increasing the spiking threshold without increasing the spiking reset. In this way, the spiking threshold, and the reset determine the cost function. It is conceivable that these variables may be learned, just as network connectivity may be learned. Alternatively, these values may be predetermined for various brain areas, depending on the computational target of each brain area.

**Tuning curves and quadratic programming.** For constant input signals, the instantaneous firing rates of the neurons will fluctuate around some mean value. Our goal is to determine this mean value. We will start with the most general case, and rewrite the instantaneous firing rates as 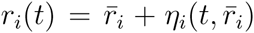, were 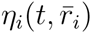 is a zero-mean ‘noise’ term that captures the fluctuations of the instantaneous firing rates around its mean value. Note that these fluctuations may depend on the neuron’s mean rate. In turn, neglecting the costs for a moment, we can average the objective function, Equation 11, over time to obtain

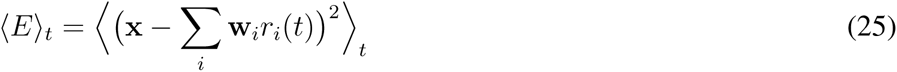

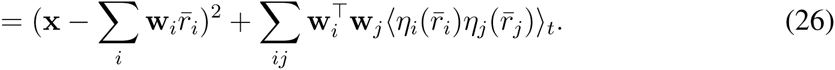

For larger networks, we can assume that the spike trains of neurons are only weakly correlated, so that 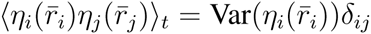. Here, 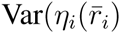 is the variance of the noise term. We obtain

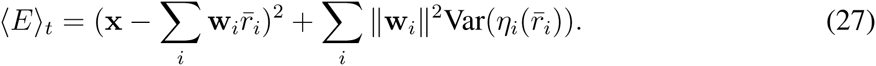

We furthermore notice that the spike train statistics are often Poisson (Boerlin et al., 2013), in which case we can make the replacement 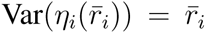. The loss function becomes then a quadratic function of the mean firing rate, which needs to be minimized under a positivity constraint on the mean firing rate. This type of problem is known as ‘quadratic programming’ in the literature. In this study, we generally focused on networks for which the contributions of the second term can be neglected, which is generally the case for sufficiently small readout weights and membrane voltage noise (see Figure S5). In this case, we obtain the multivariate version of the loss function used in Equation 4,

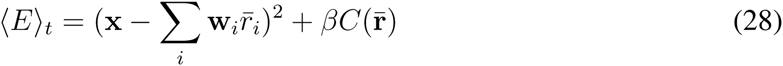

where we added the costs again. In general, quadratic programming is mathematically intractable, so the objective function must be minimized numerically. However, in networks with a small number of neurons, we can solve the problem analytically and gain some insight into the nature of quadratic programming. Here, we do this for the two neuron example (Figure 3A).

In this example, firing rates 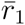 and 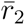 are given by the solution to the following equation:

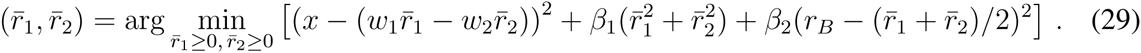

The positivity constraint partitions the solution of this equation into three regions, determined by the value of *x*: region 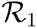 where 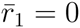 and 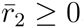, region 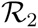 where 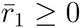 and 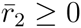, and region 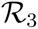 where 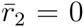 and 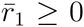 (Figure S2A). In region 2, we can then easily solve Equation 29 by setting the derivative of our loss function to zero. This gives 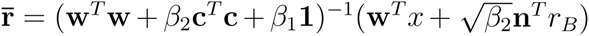 where **w** = (*w*_1_, − *w*_2_) and **c** = (1,1)/2. Looking at this equation, we see that 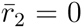 when 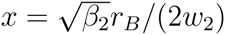 and 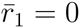 when 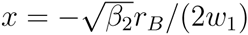. Therefore, the firing rate solution for region 2 is valid when 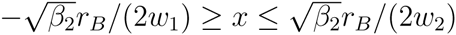. For 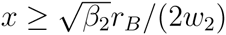, we have 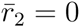 because of the positivity constraint in Equation 29. This is region 3. We can calculate 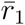 by setting 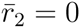 and then minimizing the loss function. This gives 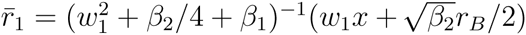. Similarly, for 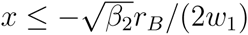 we have 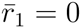 because of the positivity constraint. This is region 1 and we obtain 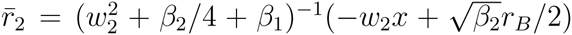. The firing rates within each region are given by a simple linear projection of 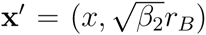, although the size and direction of this projection is different in each region. As such, the solution to this quadratic programming problem is a piece-wise linear function of *x*.

In networks with larger numbers of neurons, the solution will still be a piece-wise linear function of *x*, although there will be more regions and the firing rate solutions are more complicated because more neurons are simultaneously active. In contrast, the transformation from firing rates to *x*′ is very simple (Equation 8). It is given by a simple linear transformation, and is region independent (Figure S2B).

**Readout weights and cost terms: 1-d and 2-d example.** There are a number of free parameters in our model. These are the cost terms and the readout weights {*w_jk_*}. The choice of these values determine the precise shape of tuning curves. In general however, the precise values of these terms has little influence on the coding capability of our system, once certain properties have been satisfied, which we will outline here.

The cost term that we use for our first example (Figure 1), and for our 2-d bump shaped tuning curves (Figure 6 and Figure S7C, D) is a quadratic cost 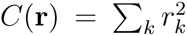. (Here, and in the following, we will no longer distinguish *r_i_*, the instantaneous firing rate used in the spiking network, and 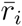, the mean firing rates used in quadratic programming. Both notations apply depending on the context.) This cost term encourages the system to find a solution in which all neurons share in the signal representation. For our homogenous monotonic tuning curve examples (Figure 3A, B and Figure S2 and S3A) we use a homogeneous quadratic cost 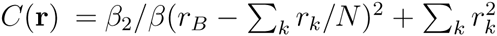, which biases the system towards a background firing rate value of *r_B_*, where *β*_2_/*β* represents the extent of this bias. For our inhomogenous monotonic tuning curve examples (Figures 2, 3C, 4, 5B, D, 6G-J and Figures S1, S3B, C and S5), we use a slightly more general *biased* quadratic cost 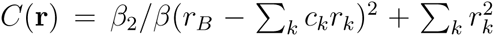. This is similar to the homogenous cost, but with a sum weighted by {*c_k_*} rather than 1/*N*. This allows us to introduce some inhomogeneity into our system. The additional bias term in this heterogeneous cost is equivalent to an additional readout dimension. To see this, consider a 1-dimensional system with 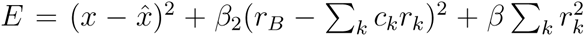. We can rewrite this as a 2-dimensional system with 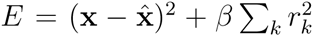, where 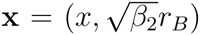 and 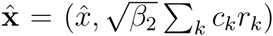. Throughout this work, our choice of cost function is motivated by our goal of making direct comparisons to data from various systems. For example, in the oculomotor system, the population firing rate is approximately constant (Aksay et al., 2000) for all values of *x*, which is a property that may facilitate constant muscle tone in the eye. We accommodate this property by choosing a biased quadratic cost term. Nonetheless, our general quadratic programming predictions about optimal compensation predictions are qualitatively similar regardless of our cost choice (Figure 4 and Figure S4).

The other important parameters that we must choose are the readout weights. For the sake of consistency and ease of comparison, we use similarly valued readout weights across several figures (Figures 2, 3C, 4, 5B, D and Figures S1, S4 and S5A, C). We have chosen values that are regularly spaced, with the addition of some random noise. Specifically, we set the readout to *w_i_* — [*w_min_* + (*w_max_ w_min_*)(*i* − 1)/(N/2 − 1)]*ξ_i_* for *i* ∈ [1, *N*/2] and *w_j_* — [*w_min_* + (*w_max_* − *W_min_*)(*j* − *N*/2 − 1)/(*N*/2 − 1)]*ξ_j_* for *j* ∈ [*N*/2 + 1, *N*]. Also, we set 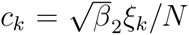 with 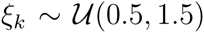(0.5,1.5). The parameter values that we use are chosen so that we can make direct comparisons between our framework and the oculomotor system (Figure 5A, B). These values are *w_min_* = 0.01 and *w_max_* = 0.05. It is interesting to observe that the tuning curves of the oculomotor system are consistent with a random readout. Indeed, we find that there are many possible parameter values that produce systems with similar performance levels (Figure S3). This suggests that the performance of the oculomotor system is adequate, even with a random readout and the resulting random (if symmetric) connectivity. For our bump-shaped tuning curve example, the parameter values that we use are plotted (Figure 6A, G).

All of our quadratic programming predictions for tuning curve shape and our optimal compensation predictions for tuning curve shape change still hold for alternative choices of readout weights (Figure 3 and Figure S3), once the following properties are satisfied. First, it is important that the readout vectors span the space of the signal that we want our network to represent. Otherwise, the system will not be able to represent signals along certain directions, or compensate for neuron death. Second, it is important to set the scale of the readout, so that the cost does not dominate. There is a natural scaling that we can use to avoid such problems. We require the size of firing rates to be independent of network size, 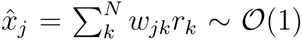, so we must have 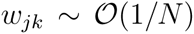. As a consequence, the off-diagonal elements of the recurrent connectivity are small 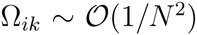, for *i* ≠ *k* (Equation 20), and if we assume that the diagonal elements are on the same order of magnitude, then 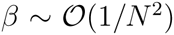. This scaling provides a principled basis for parameter choices in our model.

We may also want to scale our decoder weights without changing the shape of our tuning curve prediction (Figure S5B). To do this, the spiking cost parameter *β* and the membrane potential leak *λ* must also be scaled together. Specifically, if the readout weights are given by {*α* × *w_jk_*}, where *α* is a scaling parameter that characterizes the size of the decoder weights and {*w_jk_*} are fixed decoder weights, then the spiking cost parameter must be set to *α*^2^ × *β* and the membrane potential leak must be set to *α* × *λ*. We can see that this preserves the shape of tuning curves by looking at the resulting structure of our loss function (Equation 2):

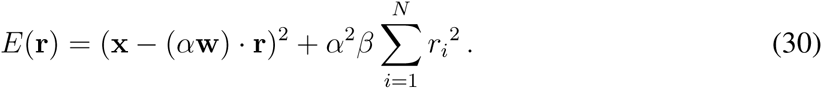

As before, the minimum of this loss function gives firing rates in units of the membrane potential leak (*αλ*). Therefore, we must divide **r** by (*αλ*) to obtain firing rates in units of Hz. Our loss function then becomes:

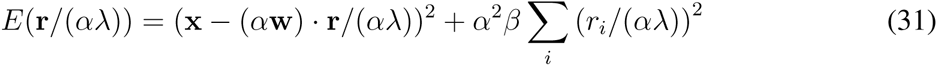

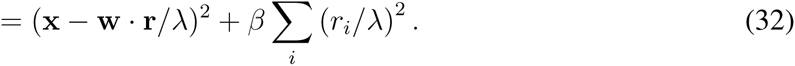

This loss function is independent of *α*, and so, using this scaling the optimal tuning curves will have the same shape for all values of *α*.

**Readout weights and cost terms: V1 example.** There are many possible choices of decoder weights {**w***_i_*} that provide a faithful representation of a signal. In positive sparse coding, we choose the decoder weights that provide the most efficient signal representation, for a sparse cost term (*C*(**r**) = ∑*_k_ r_k_*), under the constraint that firing rates must be positive. Here, we describe how we calculate these positive sparse coding weights, which we will use in several of our figures (Figures 7, 8 and Figures S6, 7).

We use the signal vector **x** = (*x*_1_,&, *x_j_*,&, *x_M_*) to denote an image patch, where each element *x_j_* represents a pixel from the image patch (Olshausen and Field, 1996). We quantify the efficiency of a sparse representation using the following loss function:

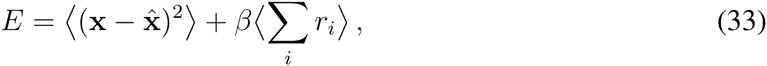

where 〈&〉 denotes an average across image patches. This is simply Equation 2 with a sparse cost term, averaged over image patches. The first term in this loss function is the image representation error. The second term quantifies the sparsity of the representation. The decoding filters that minimize this loss function will be the positive sparse coding filters for natural images.

We assume that the decoding filters are optimized to represent natural images, such as forest scenes, flowers, sky, water and other images from nature. Natural images are chosen because they are representative of the images that surround animals throughout evolution. We randomly select 2000 12 × 12 image patches from eight natural images taken from Van Hateren’s Natural Image Database (van Hateren and van der Schaaf, 1998). These images are preprocessed by removing low-order statistics, so that our sparse coding algorithm can more easily learn the higher-order statistical structure of natural images. First of all, images are centered so as to remove first order statistics:

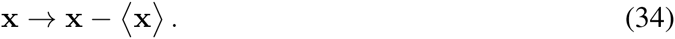

Next, images are whitened, so as to remove second-order statistics:

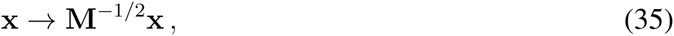

where M = 〈**xx***^T^*〉 is a decorrelating matrix.

We calculate sparse coding filters by minimizing the loss function (Equation 33) using a two step procedure. First of all, for a subset of 50 image patches, we calculate the firing rates that minimize the loss function under the constraint that firing rates must be positive:

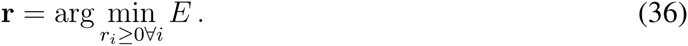

The positivity constraint reduces the representational power of the neural population, so, to counteract this, we use a large population containing twice as many neurons as signal dimensions, *N* = 2*M* and we initialize our decoder matrix to *w_jk_* = *δ_j,k_* — *δ_j,k+M_* to ensure that our neural population can easily span the signal space throughout the sparse coding calculation.

Next, we update our kernels by stochastic gradient descent on the loss function, using the optimal firing rates calculated in the previous step:

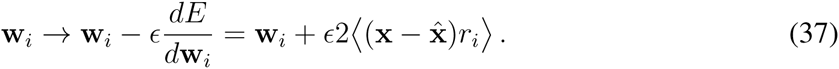

Kernel weights are normalized for each neuron, so that they do not become arbitrarily large. This calculation is similar to sparse coding calculations performed before (Olshausen and Field, 1996), except that we enforce the additional requirement that firing rates must be positive. This constraint was used before for sparse non-negative matrix factorization (Hoyer, 2004, 2003). However, this method requires an additional constraint that decoding weights be strictly positive, so that the nonnegative matrix factorization method can be applied. This additional constraint will reduce the efficiency of the representation, because every additional constraint has the property of reducing the efficiency by precluding solutions that may be more efficient, but that violate the additional constraint.

To compare the properties of optimal compensation in this positive sparse coding model to experiment, we calculate the orientation tuning of neurons in our model using a protocol similar to that used in the experimental work (Crook and Eysel, 1992). Specifically, we drive our model using an orientated edge-like stimulus. These stimuli are Gabor filters, with a profile similar to the sparse coding filters. We calculate the firing rate response of each neuron using Equation 36 for 16 different stimulus orientations. At each orientation, the Gabor filter is positioned at regularly spaced locations along a line perpendicular to the Gabor filter orientation and the maximum firing rate of each neuron along this line is taken as the firing rate response of that neuron at that orientation. These firing rate responses form the orientation tuning curves that we can compare to experimental recordings.

**The impact of neuron death.** We calculate the impact of neuron death on tuning curve shape with and without optimal compensation (Figure 4). To calculate tuning curve shape without compensation we solve Equation 4 in the intact state. We then constrain the firing rates of neurons selected for death to zero and calculate the impact of this on our signal representation. To calculate the impact of neuron death with optimal compensation, we solve Equation 5.

We also calculate the impact of neuron death and optimal compensation in an over-complete positive sparse coding model (Figure S7A). To do this, we define an over-completeness factor, *M*, given by *M* = *N/d*, where N is the number of neurons in the representation and d is the signal dimension. In the sparse coding calculation above, we had *M* = 2. We can use this to define *N*_2_ = 2*d* and 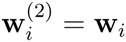, where **w***_i_* is a sparse coding readout vector. The over-complete readout vectors that we use are similar to the sparse coding readout vectors, but with some multiplicative noise (to avoid parallel readout weights). Specifically, we obtain new over complete readout vectors 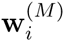 using 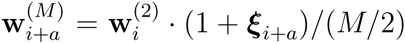, where 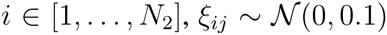, *a* = *mN*_2_ and *m* ∈ [0,&, *M*/2 − 1]. We use this method to calculate readout weights for over-complete representations because the sparse coding calculation becomes computationally prohibitive for large values of *M*.

**The impact of neuron death in a spiking network.** We investigate the impact of neuron death in our spiking model by simulating the network both before and after neuron death (Figure 2). Specifically, we use the Euler method to iterate the dynamics described by Equation 19 and we measure the readout 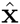, the spike trains s and the membrane potentials V. We simulate the death of a neuron by setting all the connections to and from that neuron to zero. This is equivalent to silencing a neuron. We continue to measure the network activity and the readout after neuron death.

The integrate-and-fire model that we have described can produce spikes at arbitrarily large firing rates. This can be a problem, especially when neurons die and the remaining neurons compensate with extremely high, unrealistic firing rates. To avoid this, we include a form of adaptation in our spiking model. Specifically, we extend our spiking rule so that a neuron *i* will only spike if its firing rate *f_i_* is lower than a maximum *f_max_*. Here, the firing rate f is given by the following differential equation:

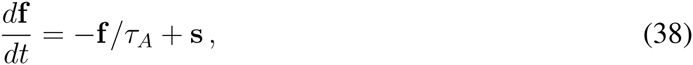

and *τ_A_* is the timescale of adaption. This timescale is much slower that the timescale of spiking dynamics *τ*.

We use this extended neural model to calculate the recovery boundary in a system that is optimized to represent a 1-d variable (Figure S1). We kill neurons in random order until the representation error averaged across signal values is more that 10% larger than the representation error in an identical system without adaptation constraints. We then average across trials to obtain a value for the recovery boundary. This working definition for the recovery boundary captures the serious degradation in representation performance that characterizes the notion of a recovery boundary.

We can also calculate the firing rates and recovery boundary of our extended integrate-and-fire model using quadratic programming (data not shown). Specifically, the firing rates of this network are given by the solution of the following optimization problem:

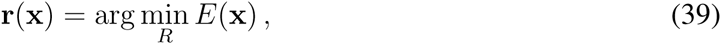

where *R* denotes the set of constraints that the firing rates must satisfy:

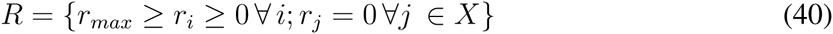

and where *X* is the set of all dead (or silenced) neurons.

**Tight balance of excitation and inhibition.** In simulations of our spiking model, we see that the balance of excitation and inhibition coincides with optimal coding (Figure 2D, G). We define the excitation *E_i_* received by a neuron as the total input current received through excitatory synapses, and inhibition *I_i_* as the total input current received through inhibitory synapses:

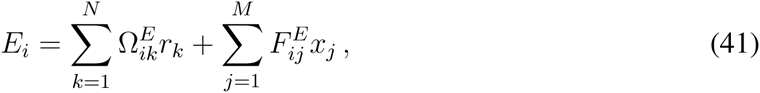

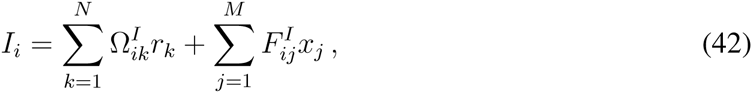

where 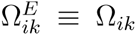 unless *Q_ik_* < 0, 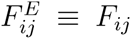 unless *F_ij_x_j_* < 0, 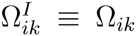 unless Q*_ik_* > 0 and 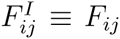 unless *F_ij_x_j_* > 0. We can understand why excitation and inhibition are balanced during optimal coding by first observing that the value of the loss function is small during optimal coding. Consequently, the membrane potential is small, because the membrane potential is a linear transformation of 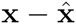 (Equation 14). Finally, for membrane potentials to be small, we observe that there must be a balance of excitation and inhibition (Equation 14, Equation 41, Equation 42). As such, the balance of excitation and inhibition is a signature of optimal coding.

We can also understand this balance using scaling arguments. As we have discussed, the recurrent connection strengths are small 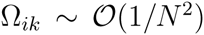. However, the spiking thresholds are also small 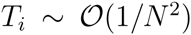, because 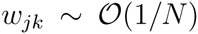 and 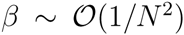 (Equation 24). Now, because recurrent input is summed across the entire population of *N* neurons, we find that 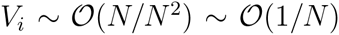. This is *N* times larger than the spiking threshold. In order for the membrane potential to be on the same order of magnitude as the spiking threshold, there must be a precise cancellation of excitatory and inhibitory inputs. This balance of excitation and inhibition is much tighter than balanced networks with random connectivity (van Vreeswijk and Sompolinsky, 1996, 1998). In randomly connected networks the balance between excitation and inhibition is of order 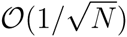 (van Vreeswijk and Sompolinsky, 1998), whereas in the tightly balanced networks that we consider, these fluctuations are 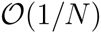.

**Optimal compensation proof.** The spiking network that we use is capable of rapidly implementing optimal compensation, without requiring any synaptic plasticity mechanisms. We can prove this by showing that an optimal network of *N* − 1 neurons is equivalent to an optimal network of *N* neurons after the death of one neuron.

For the sake of argument, we suppose that the *N^th^* neuron dies. At a mechanistic level, the death of this neuron is equivalent to cutting all the connections to and from the dead neuron and to our readout 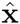. Therefore, a network where the *N^th^* neuron has died is equivalent to a network with *N* − 1 neurons with readout matrix 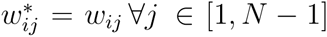 and ∀*i* ∈ [1, *M*], with feed forward connectivity 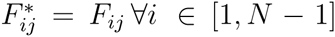 and ∀*j* ∈ [1, *M*], with recurrent connectivity 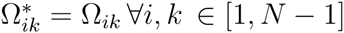 and with spiking thresholds 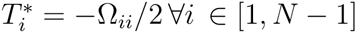.

Now, we compare this damaged network to an optimal network consisting of *N* − 1 neurons. To make a fair comparison, we assume that this network has the same readout matrix **w**′ = **w**^*^ as the reduced damaged network. Then, the recurrent connectivity for this network is given by Ω′ = Ω^*^, the feed forward connectivity is given by **F**′ = **F**^*^ and the spiking thresholds are given by **T**′ = **T**^*^. This configuration is equivalent to the reduced damaged network. Therefore, a spiking neural network who’s neurons are individually tuned to represent a signal optimally before cell death will perform optimal compensation and provide an optimal signal representation after cell death.

## Author Contributions.

D.G.T.B., S.D and C.K.M. designed the study. D.G.T.B. and C.K.M. performed the analysis. D.G.T.B and C.K.M wrote the manuscript.

## Acknowledgements.

We thank Tony Movshon, Pedro Gonçalves, and Wieland Brendel for stimulating discussions and Alfonso Renart, Claudia Feierstein, Joe Paton and Michael Orger for helpful comments on the manuscript. S.D. acknowledges the James McDonnell Foundation Award and EU grants BACS FP6-IST-027140, BIND MECT-CT-20095-024831, and ERC FP7-PREDSPIKE. C.K.M. acknowledges an Emmy-Noether grant of the Deutsche Forschungsgemeinschaft and a Chaire d’excellence of the Agence National de la Recherche.

## Supplemental Figures and Legends

**Figure S1.**
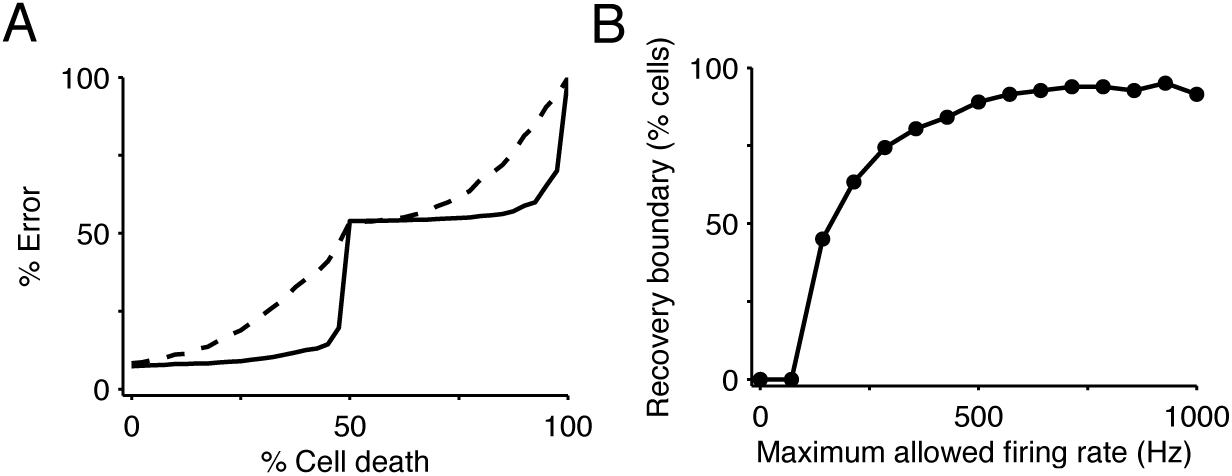
The recovery boundary (related to Figure 2) **(A)** We progressively kill neurons in our balanced network model and calculate the signal representation error (black line). The signal representation error is given by the average; loss function value (average of Equation 2), where averages are taken across signal values and time and are written as a percentage of the maximum average error. Neurons with positive read-out weights are knocked out first, followed by neurons with negative read-out weights. The initial increase in error is very small, because optimal compensation successfully recovers the signal representation. However, when neuron death hits the recovery boundary, in this case at 50% neuron death (equivalently, 100% death of neurons with positive read-out weights), there is a sudden break-down in signal representation. We also calculate the signal representation error where the maximum firing rate of each neuron is constrained to be 100 Hz (dashed line). We constrain the maximum firing rate of neurons using an adaptation mechanism (see Methods). In this case the recovery boundary occurs much earlier. **(B)** We calculate the average recovery boundary as the maximum allowed firing rate value decreases from 1000 Hz to 0 Hz. The average recovery boundary is calculated by killing neurons in random order in each trial, and then taking an average recovery boundary value across different trials. We find that the recovery boundary occurs earlier (with less neuron death) in systems with low maximum firing rate values compared to networks with high maximum firing rate values.

**Figure S2.**
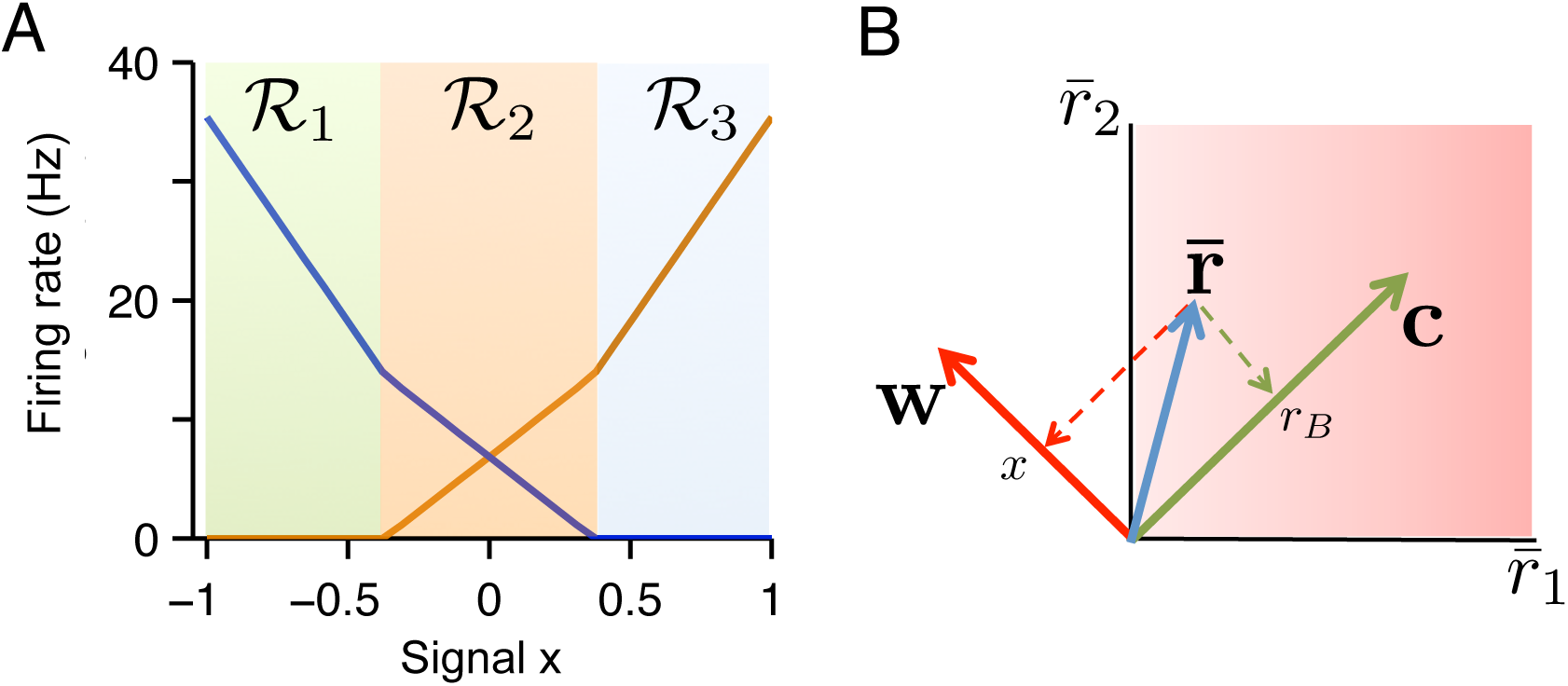
The geometry of quadratic programming, (related to Figure 3) **(A)** Tuning curves calculated for a two neuron example using Equation 29. The tuning curve solution can be decomposed into three regions: region 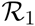 where 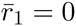 and 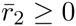, region 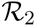 where 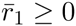 and 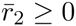, and region 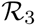 where 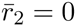 and 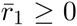. In each region the firing rate solution is given by a different linear projection of 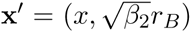, where *r_B_* is the background firing rate. **(B)** The transformation from firing rate space to signal space is given by a simple linear projection of firing rates onto the read-out vector **w** (for the signal *x*) and **c** (for the background firing rate *r_B_*). Unlike the transformation from signal space to firing rate space in **A**, this transformation is not region dependent.

**Figure S3.**
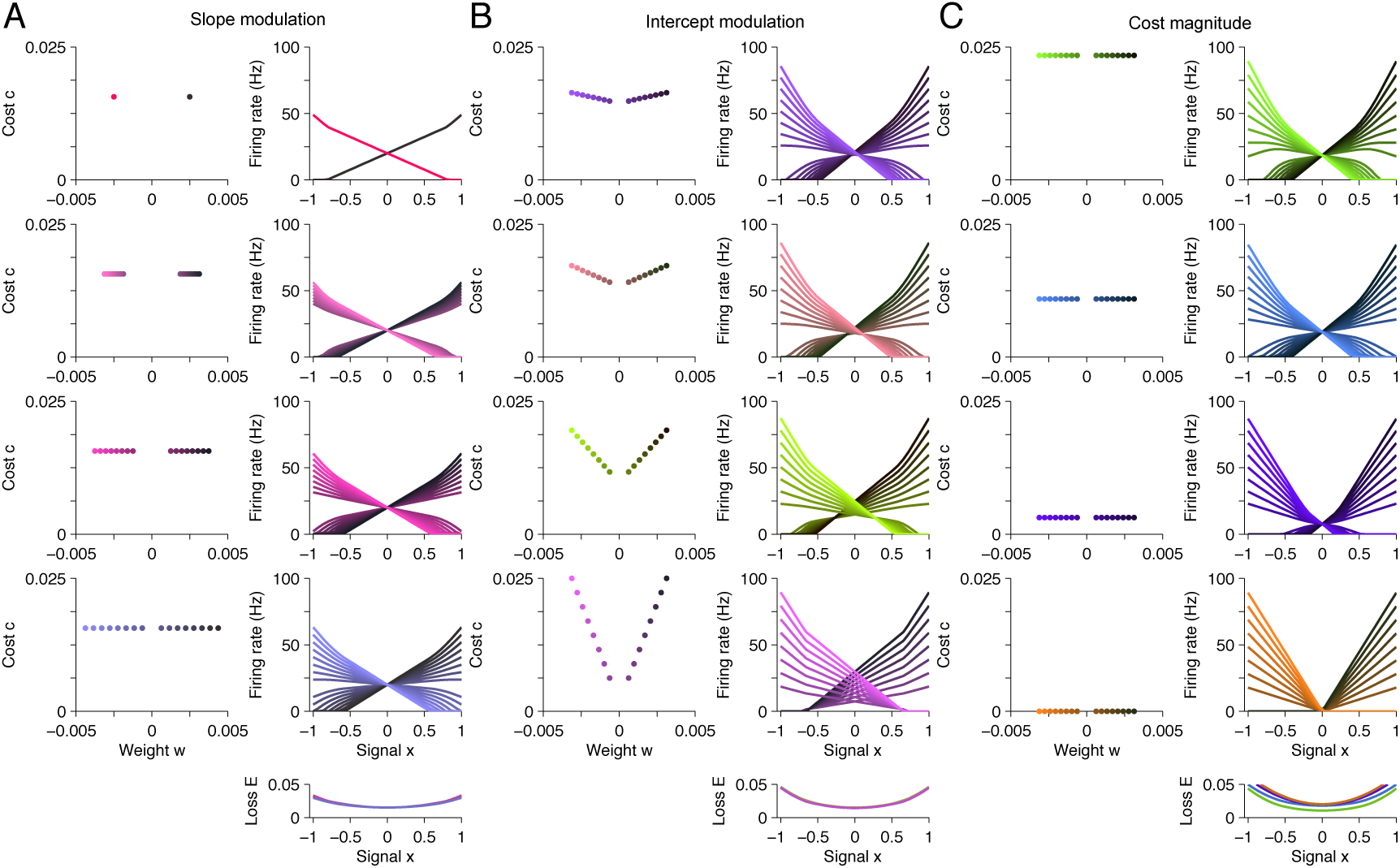
A taxonomy of tuning curve shapes (related to Figure 3). Here, we explore the relationship between tuning curve shape and the choice of parameters in our 1-d system. Specifically, we calculate tuning curve shape using quadratic programming (Equation 4) for 12 distinct systems, each with different parameter values. **(A)** The left column shows the read-out weights {*w_i_*} and cost terms {*c_i_*} for each system. The right column shows the tuning curves calculated using quadratic programming (Equation 4). Each row corresponds to a distinct neural population. In this panel, we increase the spread of read-out weights from top to bottom (left column), and we find that the range of tuning curve slopes increase (right column). The loss is approximately invariant to this modulation (right column, bottom). **(B)** Similar to **A**, except that we modulate the cost terms from top to bottom (left column), and we observe that the range of tuning curve intercept values increase (right column). Again, the loss is approximately invariant to this modulation (right column, bottom). **(C)** Similar to **A** and **B** except that this time we reduce the magnitude of the cost terms from top to bottom (left column), and we observe that the intercept values decrease, until tuning curves do not overlap with each other (right column). This time, the loss increases marginally (right column, bottom). This happens because the system is unable to represent the background task variable (see Methods). In this figure, each system contain 16 neurons (some tuning curves overlap in **A**) and the biased quadratic cost term as in all previous 1-d examples.

**Figure S4.**
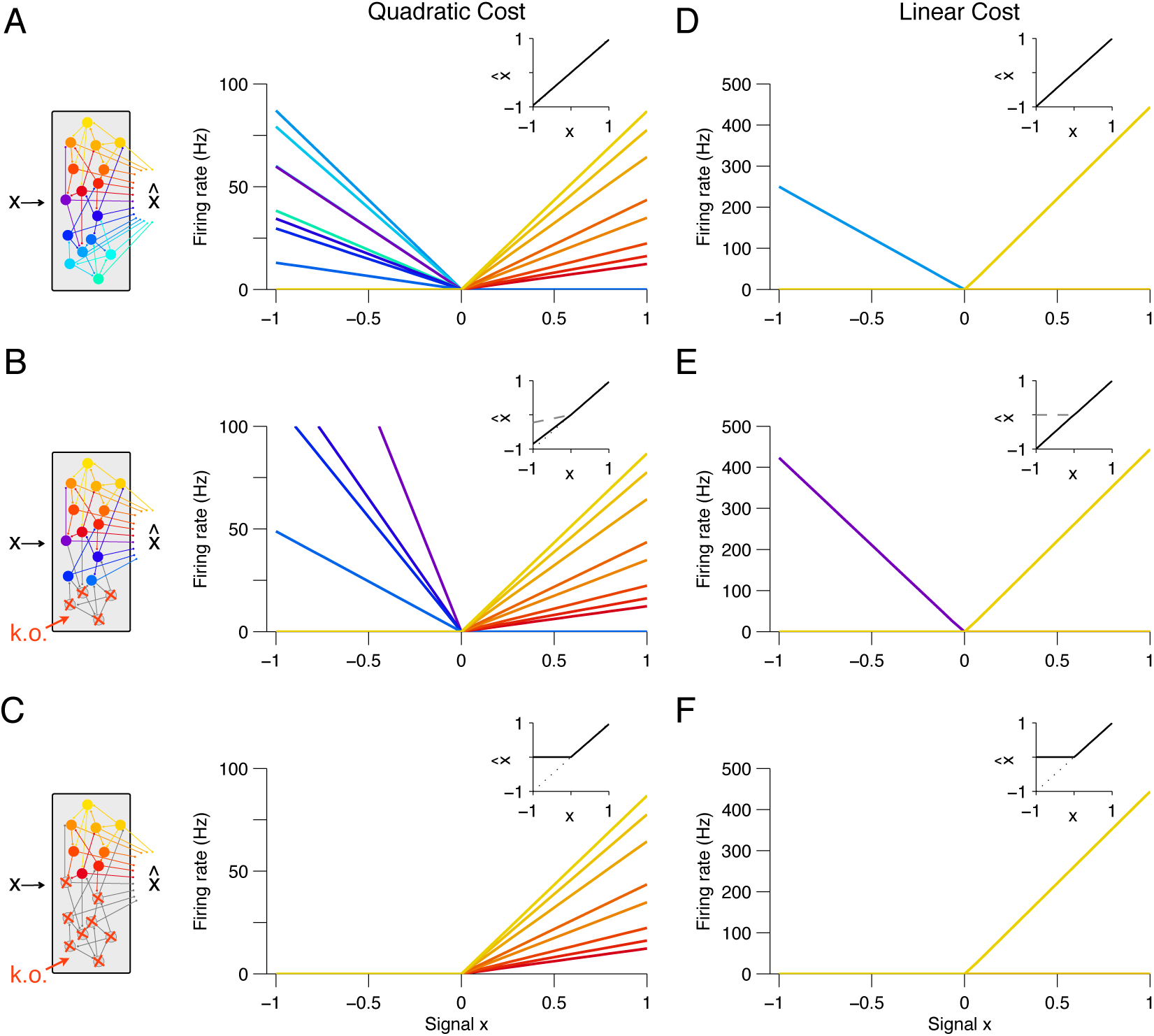
Optimal Compensation in systems with a linear cost or a quadratic cost (related to Figure 3). **(A)** Optimal tuning curves calculated using quadratic programming (Equation 4), but with a quadratic cost term, instead of the biased quadratic cost term. In this system, tuning curves do not overlap, because there is no term to bias the firing rates towards a background task variable *r_B_*. Therefore, this is a poor model of the oculomotor system (Figure 5a). **(B)** We knock-out four neurons with negative read-out weights and calculate tuning curves following optimal compensation using Equation 5. As before, the slopes of similarly tuned neurons increase to compensate. **(C)** When all the neurons with negative read-out weights are knocked out, optimal compensation is unable to recover signal representation (inset), because the system has crossed the recovery boundary. **(D-F)** Similar to **A–C**, but for a system with a sparse coding linear cost instead of a quadratic cost. **(D)** In this sparse representation, all neurons are silent, except for the neuron with the largest positive read-out weight and the neuron with the largest magnitude negative read-out weight. These two neurons represent the entire signal, because they can do so with the smallest firing rates, and hence, with the smallest cost. **(E)** We knock-out four neurons with negative read-out weights, including the neuron that was active in **D**, and we calculate tuning curves following optimal compensation. A single neuron compensates for neuron death. This is the neuron with the largest magnitude negative read-out weight (of the remaining neurons). Note that signal representation without optimal compensation is especially poor compared to the system with optimal compensation (inset). **(F)** When all the neurons with negative read-out weights are knocked out, optimal compensation is unable to recover signal representation.

**Figure S5.**
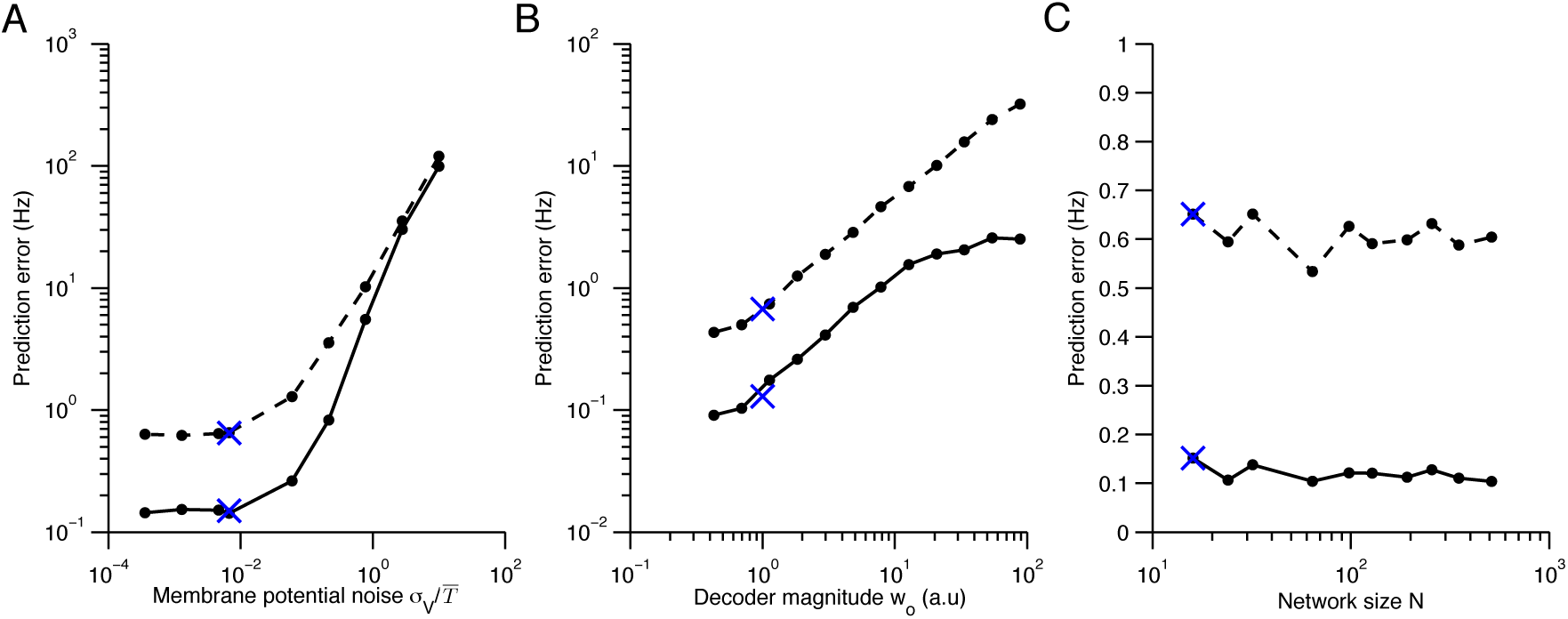
Quadratic programming firing rate predictions compared to spiking network measurements. **(A)** The impact of membrane potential noise *σ_V_* on firing rate predictions is calculated by comparing quadratic programming predictions and firing rate measurements in network simulations. We calculate the prediction error of quadratic programming by taking the average absolute difference between quadratic programming predictions and spiking network measurements (solid line) and by taking the standard deviation of these measurements about the predictions (dashed line). Averages are calculated across neurons, stimuli and time, using networks with tuning curve shapes similar to previous figures (Figure 4A). In the parameter range used throughout this paper (indicated by a blue cross), the prediction error is small and robust to changes in noise. As expected, the error increases when the noise is the same order of magnitude as the mean spiking threshold, 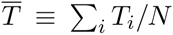 **(B)** Similarly, the prediction error increases as the size of the decoder weights increase. Here, α is a scaling parameter that characterizes the magnitude of the decoder weights (see Methods). We find that the error is small around the parameter range used throughout this paper (*α* = 1). However, for larger values of *α* the quadratic programming prediction degrades. Again, this is expected because *α* determines the resolution with which our spiking network can represent a signal. As such, our predictions are most accurate in the very regime that we are most interested in - where optimal coding is possible. Note that we must also scale the spiking cost and the membrane potential leak with *α* so that the shape of tuning curves are preserved, allowing for a fair comparison across decoder scales (see Methods). **(C)** The network size *N* has little influence on the prediction error. Again, read-out weights and cost parameters are scaled so that tuning curve shape is invariant to changes in *N* (see Methods).

**Figure S6.**
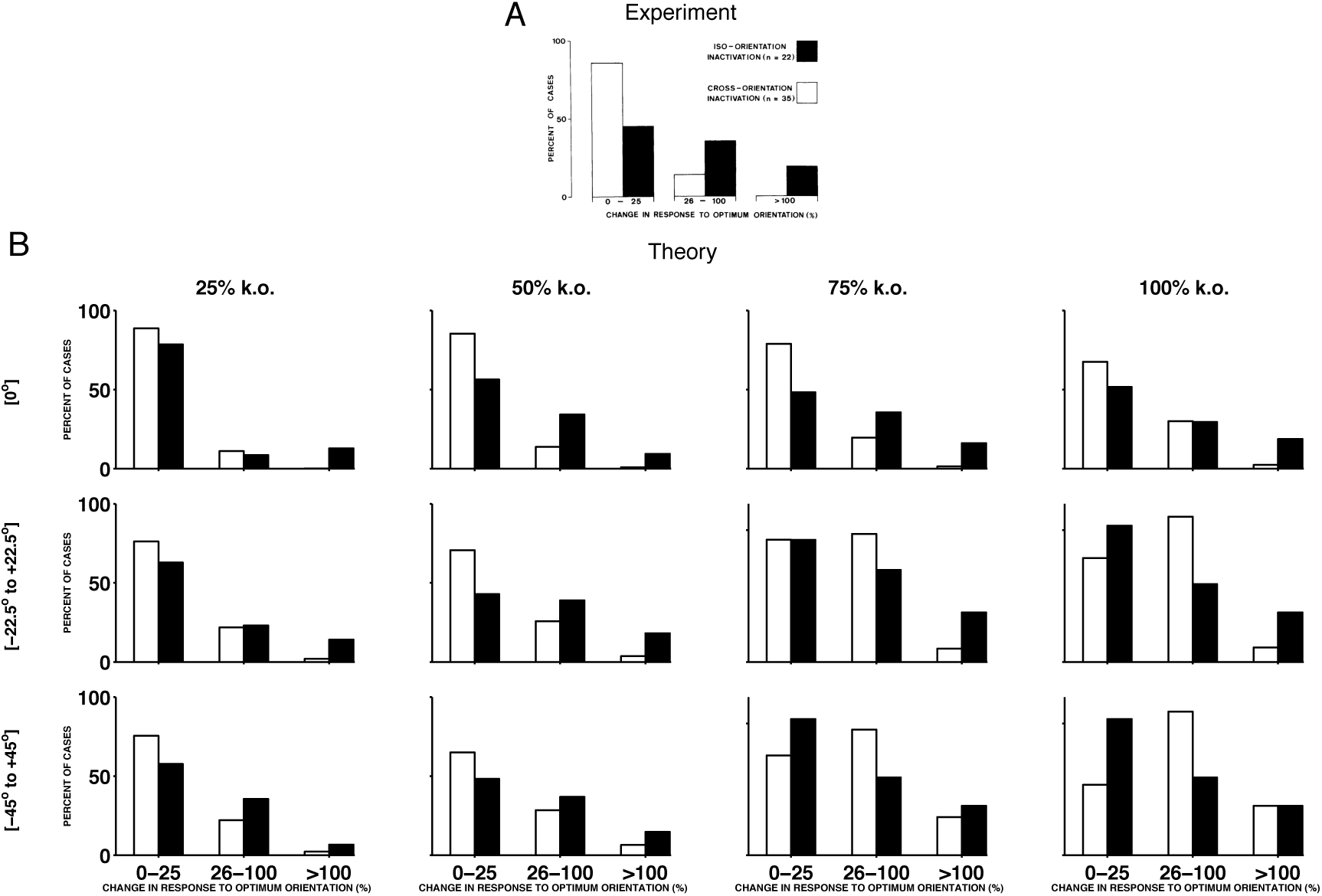
Histograms of experimental responses to neuron silencing in V1, compared to theoretical predictions using a range of different parameters. **(A)** Histogram of changes in the firing rate at preferred stimulus orientations following GABA-ergic silencing. Firing rate change for neurons with tuning preferences that are similar to the recorded neurons (iso-orientation inactivation) are counted separately to changes in neurons with different tuning preferences (cross-orientation inactivation). These results are reproduced from Crook and Eysel (1992). **(B)** Histograms of preferred compensation firing rate changes in positive sparse coding neurons, again with iso-orientation and cross-orientation neurons counted separately. Each histogram corresponds to the theoretical prediction obtained by knocking out different percentages of neurons (25%, 50%, 75% and 100%), across different ranges of preferred orientations (at 0*°* only, from −22.5*°* to 22.5*°*, and from −45*°* to 45*°*). We explore the full parameter space of our model, because the exact amount of neuron death in the experiments from Crook and Eysel (1992) is unknown. We find that when 50% or 75% of neurons with preferred orientations at 0*°* are knocked out, the form of the predicted histogram is similar to the experimentally recorded histogram, with a greater proportion of iso-orientation inactivations having an impact on the firing rate at the preferred stimulus orientation, compared to cross-orientation inactivation. In these calculations, we use the same sparse coding model for each histogram, with four times as many neurons as stimulus dimensions.

**Figure S7.**
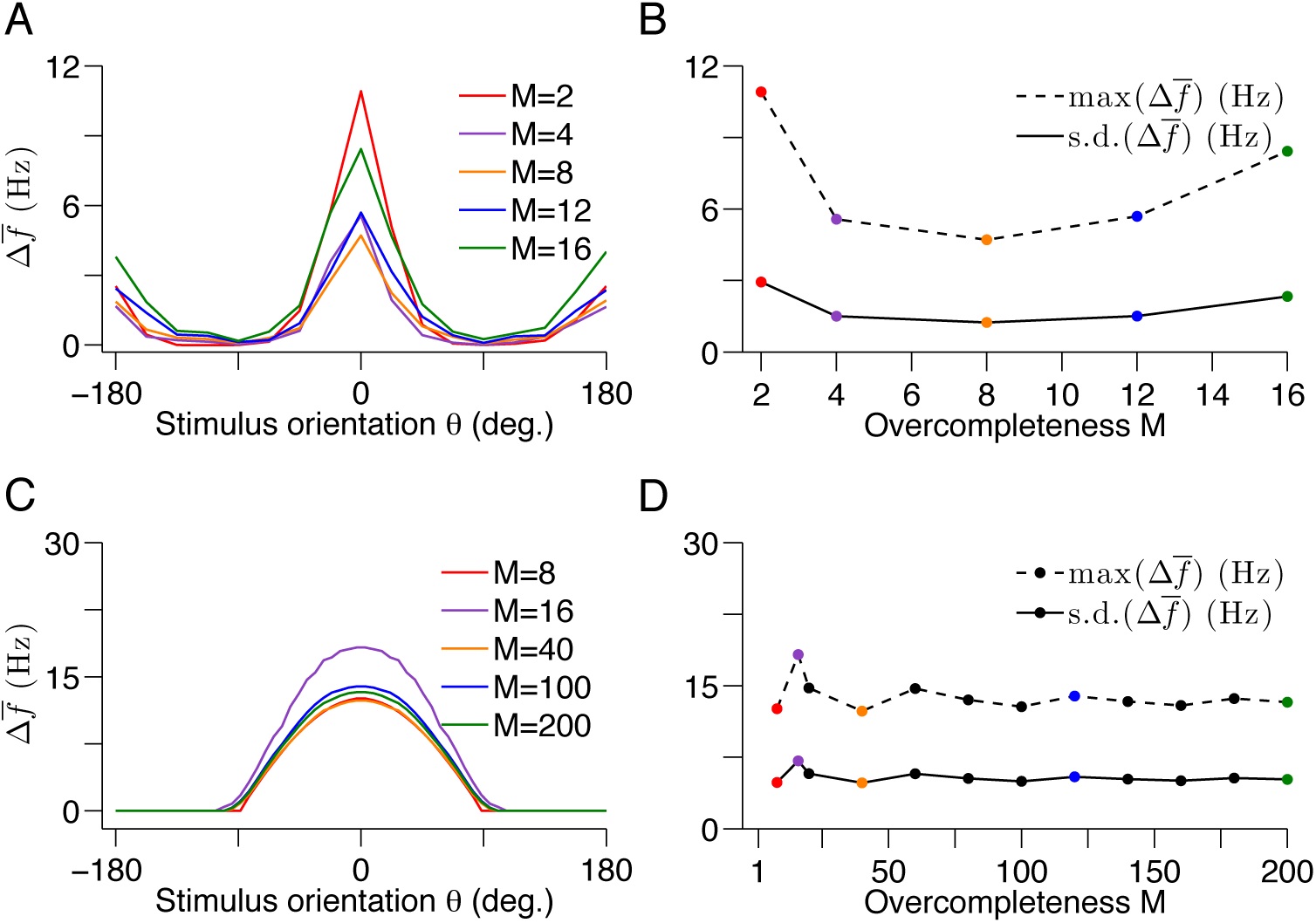
Optimal compensation in V1 models with different degrees of over-completeness (related to Figure 8) **(A)** The visual cortex contains many more neurons than input dimensions. To investigate the impact of this over-completeness, we calculate the average change in tuning curve shape following optimal compensation in our sparse coding model of V1 for increasing degrees of over completeness (see Methods). Here, the over completeness factor, *M* is given by *M* = *N/d,* where **N** is the number of neurons and d is the signal dimension. The form of the tuning curve changes is unaffected by the degree of over-completeness, but there are some fluctuations in the overall change. **(B)** As the degree of over completeness *M* increases, the average change fluctuates moderately. These fluctuations are the result of inhomogeneities in our V1 model, which have a larger effect when the over-completeness factor is small. **(C)** Similar to **A**, but for the 2-d bump-shaped tuning curve model. We use the same model as before (Figure 6 G), but with a sparse linear cost instead of a quadratic cost. For each value of *M*, we choose decoder weights that are evenly spaced on the unit circle. This produces evenly spaced bump-shaped tuning curves. We knock out neurons in this model and calculate the average change in tuning curve. In this case, we can easily calculate the impact of optimal compensation for systems with high degrees of over-completeness, such as the visual cortex because the dimensionality of the problem is much lower. **(D)** The impact of optimal compensation fluctuates moderately for low values of *M*, as in the sparse coding model. However, as the degree of over completeness increases, the average change in tuning curve shape converges, as the fluctuations average out. The maximum and standard deviation of the average tuning curve change are calculated across all stimulus orientations. In both the full sparse coding model and the 2-d model, we knock out neurons with preferred orientations between −22.5*°* and 22.5*°*.

